# *Drosophila* host defense mechanisms against filamentous fungal pathogens with diverse lifestyles

**DOI:** 10.1101/2025.08.20.671119

**Authors:** Guiqing Liu, Yao Tian, Mark Austin Hanson, Prince Kumar Sah, Jun Li, Bruno Lemaitre

## Abstract

Entomopathogenic fungi serve as powerful regulators of insect populations in nature. However, how immune effectors combat fungal pathogens remains incompletely understood. We employ *Drosophila melanogaster* as a genetically tractable model to dissect immune defense mechanisms against diverse fungal pathogens. We show that the Toll pathway is the key determinant of immunity against all species tested regardless of their ecological strategy, primarily through resistance mechanisms that limit fungal proliferation. In addition, melanization, but not phagocytosis or the Imd pathway, also has a role in limiting fungal entry and proliferation. Additionally, we show that fungal protease detection by Persephone has a quantitatively more critical role than the glucan sensor GNBP3 in the activation of the Toll pathway upon fungal infection. Our study also reveals that the fly-obligate fungus *Entomophthora muscae* employs a vegetative development strategy to hide from the host immune response. These findings reveal that *Drosophila* immune mechanisms effectively defend against a broad range of fungal pathogens, while highlighting striking adaptations to overcome these defenses in highly specialized fungal pathogens such as *E. muscae*.

## Introduction

Fungal infections are widespread in nature and exert a profound impact across a wide range of organisms, including agriculturally significant plants, insects, and humans (Brown, 2023; Dean et al., 2012; Wang and Wang, 2017). Interactions between fungal pathogens and their hosts are highly dynamic and intricate. This complexity arises from the vast evolutionary diversity of fungal pathogens and the ever-changing host microenvironments that influence their infection (Brown, 2023; Dean et al., 2012; Wang and Wang, 2017). An enhanced understanding of host-fungus interactions could lead to the development of more effective strategies for managing fungal infections of global importance in agriculture, public health or environmental management (Casalini et al., 2024; Rommelaere et al., 2025).

Fungal pathogens have evolved within different niches. Saprotrophic fungi such as *Cryptococcus neoformans* and *Aspergillus fumigatus* primarily inhabit the environment but have evolved independent mechanisms that facilitate animal infections (Morelli et al., 2021; Ristow et al., 2023). In contrast, other fungal pathogens, including *Candida albicans* and entomopathogenic fungi such as *Metarhizium* and *Beauveria* species, have co-evolved with their hosts (Brown, 2023; Hu et al., 2014). Some, under strong selection to exploit their hosts for survival, have even become obligate pathogens as seen in *Entomophthora* and *Ophiocordyceps* species (Ballou et al., 2016; De Fine Licht et al., 2017; Liu et al., 2019; Wang and Wang, 2017). Unlike human-pathogenic fungi, entomopathogenic fungi can hardly grow or even survive at 37°C (Siscar-Lewin et al., 2022). The genus *Metarhizium* includes generalist species capable of infecting hundreds of insect species across various orders (e.g. *M. robertsii*, formerly classified as *M. anisopliae*) and specialist species with a narrow host range (e.g. *M. acrium* and *M. rileyi*). Comparative genomic analyses suggest that generalist species evolved from specialists by broadening their repertoire of virulence factors (Gao et al., 2011; Hu et al., 2014). Intriguingly, some specialists such as *E. muscae* and *O. unilateralis* manipulate host behavior. They prompt infected insects to climb to elevated positions, affix themselves with their proboscides, and then splay their wings before dying, thereby facilitating fungal spore dissemination (Elya et al., 2018; Hughes et al., 2011).

Entomopathogenic fungi play a crucial role in regulating insect populations in natural ecosystems. Among them, the genetics and infection biology of *Metarhizium* and *Beauveria* species have been extensively studied and several strains have been developed as environmentally sustainable mycoinsecticides (Wang and Wang, 2017). Unlike bacterial and viral pathogens, which primarily utilize oral or respiratory routes, entomopathogenic fungi primarily infect insects through direct penetration of the cuticle and proliferate within the hemolymph. As a result, they must overcome multiple defenses, including cuticular barriers and both localized and systemic immune responses (Levitin and Whiteway, 2008; Mpamhanga and Kounatidis, 2024; Westlake et al., 2024).

On the pathogen side of host-pathogen interactions, entomopathogenic fungi have evolved various virulence factors to overcome or evade host defense (Cen et al., 2017; Ma et al., 2024; Qu and Wang, 2018; Tang et al., 2025; Valero-Jiménez et al., 2016; Wang et al., 2023). Comparative genomic studies indicate that generalist fungi tend to have an expanded repertoire of diverse protein families compared to specialist fungi, including cuticle-degrading proteins (proteases, carbohydrate esterase) and pathogen-host interaction proteins (Hu et al., 2014; Shang et al., 2016; Wichadakul et al., 2015). However, the pathogenic mechanisms of less-studied fungi, such as *M. rileyi* and *E. muscae*, as well as the host defense responses they elicit, remain largely unexplored (Boucias et al., 2016; Elya and De Fine Licht, 2021).

The fruit fly *Drosophila melanogaster* serves as a powerful model organism for investigating multiple defense mechanisms against fungal infections (Levitin and Whiteway, 2008; Mpamhanga and Kounatidis, 2024; Westlake et al., 2024). A range of immune receptors detect microbial molecules and damage-associated signals, triggering key immune responses such as the Imd and Toll pathways, phagocytosis and melanization. The Toll and Imd pathways coordinate the production of immune effectors by the fat body during systemic infection (De Gregorio, 2002; Rommelaere et al., 2024; Uttenweiler-Joseph et al., 1998; Valanne et al., 2011; Westlake et al., 2024). The Imd pathway controls the production of antibacterial peptide (e.g. Diptericin, Cecropin) and is critical to resist infection by Gram-negative bacteria (Carboni et al., 2022; Hanson et al., 2019). The Toll signaling pathway is critical to resist fungal infection and infection with Gram positive bacteria (Lemaitre et al., 1996; Ryckebusch et al., 2025). It can be activated by two mechanisms: if a key fungal cell wall component, *β*-1,3-glucan, binds to the secreted pattern-recognition receptor GNBP3, or if fungal proteases activate a complex extracellular cascade of serine proteases, the Toll-PO cascade. Both mechanisms converge to the activating cleavage of the Toll ligand Spätzle (Spz) (El Chamy et al., 2008; Gottar et al., 2006; Westlake et al., 2024). Binding of Spz to the Toll receptor initiates an intracellular cascade leading to the production of a battery of host defense peptides (Weber et al., 2003). Among them, genetic and *in vitro* studies have revealed the role of Drosomycin, Metchnikowin, Bomanins, Daisho and Baramicin to combat fungal infection (Clemmons et al., 2015; Cohen et al., 2020; Fehlbaum et al., 1994; Hanson et al., 2021; Levashina et al., 1995). These host defense peptides can function additively or synergistically to combat pathogens, but they can also display high specificity toward a pathogens (Hanson, 2024; Lazzaro et al., 2020). The Toll-PO cascade also activates the melanization reaction. It leads to the maturation of ProPhenoloxidases (PPO) into Phenoloxidases (PO) that produce melanin and reactive oxygen species with antimicrobial activity (Dudzic et al., 2019; Shan et al., 2023; Westlake et al., 2024). The melanization reaction and phagocytosis have been shown to contribute to surviving infection, including by fungi (Binggeli et al., 2014; Feldman et al., 2019; Lohse and Johnson, 2008; Quintin et al., 2013).

Most research on entomopathogenic fungi have focused on a few model species or genera, leaving the full extent of their diversity largely unexplored. Consequently, the roles of immune pathways and effectors in mediating host defense against different types of fungi remain poorly understood. In this study, we use *D. melanogaster* to systematically assess the roles of immune modules, pathways, and immune effectors in surviving fungal infections with varying lifestyles. By employing genetically engineered flies with single and multiple mutations in immune genes, we aim to determine: (i) which immune pathways are crucial and whether the same pathways are used against different fungal species, (ii) the relative importance of immune modules depending on the infection route, (iii) whether immune pathways function independently or synergistically to combat different fungi, and (iv) how antifungal peptides contribute individually or collectively to host defense against fungal infections.

## Results

### Fungal strains, modes of infection and immune deficient mutants

We aimed to determine whether differences in *Drosophila* antifungal defense could be revealed by using fungal pathogens with distinct lifestyles. We selected five fungal species: *Aspergillus fumigatus*, *Beauveria bassiana*, *Metarhizium anisopliae*, *Metarhizium rileyi*, and *Entomophthora muscae*. *A. fumigatus* is an opportunistic pathogen with limited ability to penetrate the insect cuticle, yet it is pathogenic when introduced systemically (Lemaitre et al., 1996). *B. bassiana*, *M. anisopliae*, *M. rileyi,* and *E. muscae* are entomopathogenic fungi capable of infecting insects through the cuticle (Elya et al., 2018; Wang and Wang, 2017). While *B. bassiana* and *M. anisopliae* have broad host ranges, *M. rileyi* predominantly infects noctuid moths with a more restricted host range (Boucias et al., 2016) and *E. muscae* is a fly-obligate pathogen (Edwards and De Fine Licht, 2024).

Spores (conidia) from *A. fumigatus, B. bassiana*, *M. anisopliae* and *M. rileyi,* can be produced *in vitro*, allowing us to use two infection methods: natural (topical) infection via immersion of flies in a spore suspension, and systemic infection by direct pricking with a needle dipped in the spore solution. In contrast, *E. muscae* spores cannot be cultured outside its host. For this species, natural infections were initiated by spore shower from cadavers of previously infected individuals as described previously (Edwards and De Fine Licht, 2024; Elya et al., 2018) and protoplasts, which can be grown *in vitro*, were used for injections. Fly stocks were maintained at 25°C, but infections were carried out at 29°C for *A. fumigatus*, *B. bassian*a, *M. anisopliae*, and *M. rileyi,* and 21°C for *E. muscae*. The immune pathway mutants and antimicrobial peptide-deficient strains used in this study have been described previously (Hanson et al., 2019; Ryckebusch et al., 2025). Most mutations were isogenized in *w^1118^* DrosDel background by seven rounds of successive backcrosses (referred to as “iso”), although some non-isogenized mutants were also included in this study (see full list of genotypes in Table S1).

### Toll and melanization contribute to surviving fungal natural infections

Survival analysis upon topical natural infection with the five fungi were performed with male *w^1118^ Drosdel flies (hereafter wild-type flies) and* flies specifically deficient for Toll (spätzle, *spz^rm7^*), Imd (Relish, *Rel^E20^*), phagocytosis (*NimC1^1^; Eater^1^*) and melanization (*PPO1 ^Δ^, PPO2 ^Δ^*), as well as compound mutant flies simultaneously lacking these four modules (*Hayan-psh^Def^; NimC1^1^; eater^1^, Rel^E20^* hereafter named *ΔIPTM* for *ΔIMD, ΔTOLL, ΔPhag, ΔMel*) (Ryckebusch et al., 2025). Survival upon natural infection reveals the major contribution of the Toll pathway, and to a lesser extent melanization to host defense against these five fungal pathogens, while deficiency in IMD and Phagocytosis have no significant effect (Figs. 1A-E). The absence of major role of the cellular response is supported by our observation that hemoless adults that have reduced number of plasmatocytes due to the overexpression of *Bax* show no or only a mild susceptibility to *B. bassiana* or *M. anisopliae*, respectively (Figs. S1A-D). The susceptibility of *spz^rm7^* flies was similar to that of *ΔIPTM* flies lacking all four immune modules indicating that Toll is by far the most important pathway to survive all the tested fungi (Table S3). However, *ΔIPTM* flies were more susceptible than *spz^rm7^* flies to *A. fumigatus,* but this likely reflects the already short lifespan of *ΔIPTM* flies (Fig. 1F), as natural infection with *A. fumigatus* did not cause a strong reduction in lifespan (Ryckebusch et al., 2025). Consistent with the findings in males, a similar role of the Toll pathway and melanization in surviving fungal infection was also observed in females, though with a lesser role for melanization in defense against *B. bassiana* in females (Figs. S2A-E).

**Figure 1.**
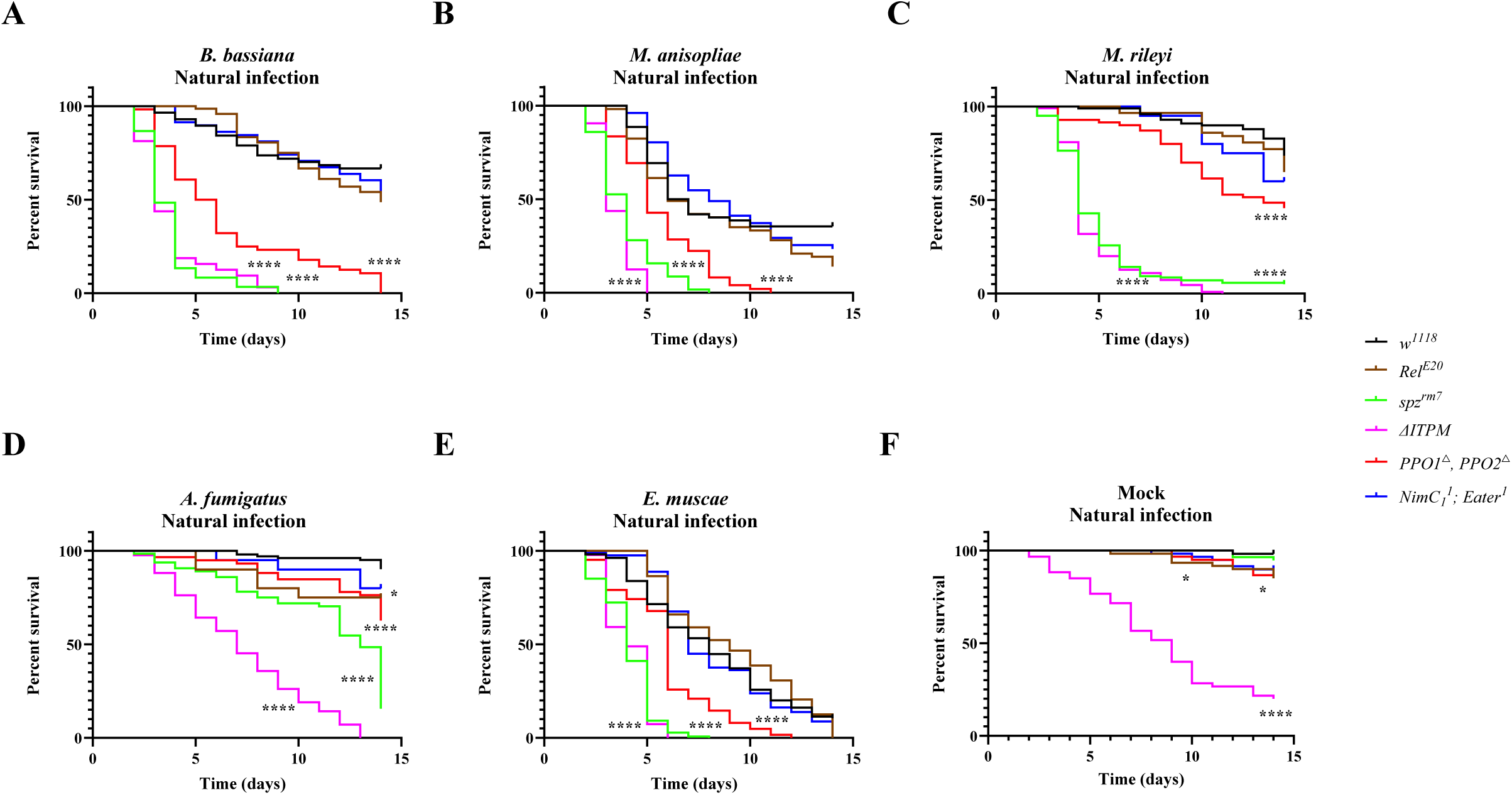
Contribution of the four main immune modules to survival upon natural infection with the five fungi. (A-E) Survival rate of male wild-type and deficient flies following infection with the five different fungi shows that *spz^rm7^* flies (P<0.0001) and *PPO1^△^, PPO2^△^* flies (P<0.0001) are severely affected in their capacity to survive infection with the five fungi compared to wild-type flies. (F) Flies that are devoid of all the four immune modules (*△ITPM*) have a significantly shorter lifespan (χ²=78.50, P<0.0001). Insects treated with 0.05% Tween 80 were used as mock controls. Data were analyzed by Log-rank test and values are pooled from at least three independent experiments. Full statistical details are available on Table S3.

The four entomopathogenic fungi were more pathogenic to *Drosophila* than the opportunistic pathogen *A. fumigatus* (Figs. 1A- E), reflecting their ability to naturally infect flies. Although *E. muscae* is a specialist fungus that exclusively parasitizes flies, it displayed the same level of pathogenicity as generalist entomopathogenic fungi *B. bassiana* and *M. anisopliae*, and the parasitic noctuid specialist *M. rileyi* to *spz^rm7^* flies (Compare 1E to Figs. 1A-C). We estimate the quantity of spores released from twenty cadavers for 24 h infection to be around 4.3×10^3^ (Fig. S2F), a much lower spore concentration than the ones used for natural infection with the other four fungi.

Collectively, our study shows that the Toll pathway is the main host defense mechanism upon natural infection with the five fungal species. Beyond the Toll pathway, melanization but not the Imd pathway or phagocytosis plays an important role. Interestingly, survival analysis shows the high pathogenicity of *E. muscae* to *Drosophila*.

### Reduced role of melanization in host defense against fungal infection upon septic injury

Injuries allow microbes to bypass the mechanical barrier normally presented by the cuticle and to directly access the nutrient-rich hemolymph. We therefore analyzed the role of the four immune modules in host defense against fungal pathogens using a septic injury mode of infection. It should be noted that *PPO1^△^, PPO2^△^* flies showed a reduced capacity to survive wounding caused by a clean injury compared with wild-type *w^1118^* flies (Fig. 2F) (Binggeli et al., 2014). Survival upon septic infection with spores of *B. bassiana*, *M. anisopliae*, *M. rileyi* or *A. fumigatus* revealed faster kinetics of killing as expected from the direct mode of infection that bypasses the crossing of the cuticle. Consistent with the observations done using a natural mode of infection, the Toll pathway was the most effective module in surviving infection by *B. bassiana*, *M. anisopliae*, *M. rileyi* and *A. fumigatus*. Interestingly, the contribution of melanization was less marked upon systemic infection compared to natural infection (compare Figs. 2A-D to Figs 1A-D). These results suggest that melanization has a key role early in the natural infection process, possibly by inhibiting fungi at the entry sites through the deposition of melanin. Unexpectedly, we observed that *Rel^E20^* flies were highly susceptible to *A. fumigatus* upon septic infection (Fig. 2D, Table S3), but not by natural infection (Fig. 1D, Table S3). This points to a role of the humoral Imd pathway against this fungus that becomes important only if the fungus can breach the cuticle.

**Figure 2.**
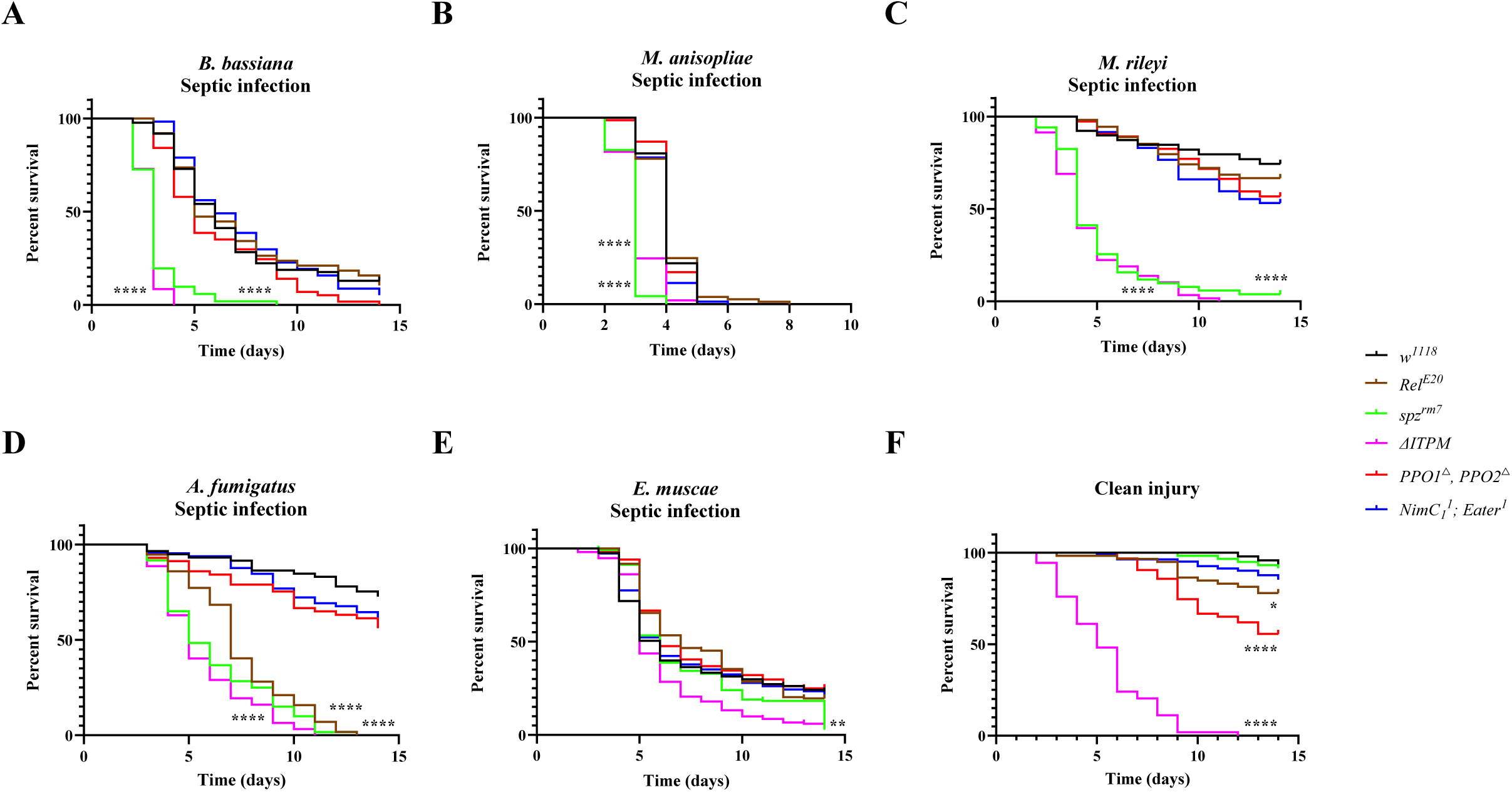
Contribution of the four main immune modules to survival upon septic infection with the five fungi. (**A-C**) Survival rate of male wild-type and deficient flies following pricking with spore suspensions show that *spz^rm7^* flies (P<0.0001) contribute most to surviving infection by the *B. bassiana*, *M. anisopliae*, and *M. rileyi* as that of natural infection. (**D**) Survival rate of male wild-type and deficient flies following pricking with spore suspensions show that *spz^rm7^* and to a lower extent *Rel^E20^* flies exhibited a strong susceptibility to infection with *A. fumigatus* (P<0.0001). (**E**) No significant difference in survival of the four main modules upon infection with *E. muscae* by the injection of protoplasts compared to wild-type flies. (**F**) Survival rate of male flies following injury with a clean needle show that *PPO1^△^, PPO2^△^* flies are severely affected in their capacity to survive wounding compared to wild-type flies (χ²=20.30, P<0.0001). There was no significant difference in survival of *△ITPM* flies upon injection with *E. muscae* protoplasts and injury with a clean needle (χ²=2.045, P=0.1527). Data were analyzed by Log-rank test and values are pooled from at least three independent experiments. Full statistical details are available on Table S3.

To analyze the host defense against *E. muscae* upon systemic infection, we injected protoplasts in wild-type and immune deficient flies. Faster killing by *E. muscae* was observed when injected with higher concentration of protoplasts (Fig. S3F), which indicates that injection of protoplasts is a valid method to initiate infection in flies. Intriguingly, flies deficient for each of the four main immune modules showed similar susceptibility to the wild-type flies after injection with *E. muscae* protoplasts. Although *ΔIPTM* flies display a significantly increased susceptibility upon injection with *E. musca* protoplasts (Fig. 2E), the difference between *△ITPM* flies injected with *E. muscae* protoplasts and clean injury was not significant (Fig. 2F, Table S3), indicating this difference rather reflects the baseline response of *△ITPM* flies to injury. Similar findings were consistently observed in females (Figs. S3A-E).

We conclude that the mode of infection alters the importance and involvement of different immune modules. Natural infection by depositing spores on the cuticle requires the melanization response to some extent, while direct injection of spores bypasses this melanization-mediated defense. On the other hand, the humoral Imd pathway plays little to no role in defense against natural infection, but an important role can be revealed specifically upon *A. fumigatus* systemic infection. We also observed that injection of *E. muscae* protoplasts kills immune mutant and wild-type flies with similar kinetics, despite immunity being involved in defense against natural infection.

### Natural infection with *E. muscae* barely induces the Toll pathway

Having revealed its contribution to combat infection with the five fungi, we next monitored the activity of the Toll pathway during the natural infection using the *Drosomycin* (*Drs*) as a read-out, a Toll-regulated gene coding for an antifungal peptide. Natural infection with the *B. bassiana* and *M. anisopliae* triggered a relatively strong expression of the Toll pathway, representing about 20-30% of the level observed 24 h after systemic injury with the model, Toll-eliciting bacterium *M. luteus* (Figs. 3A-B compared to 3F). Natural infection with *M. rileyi* triggers a significant Toll pathway activity although only at the time point of 120 h, suggesting a delayed activation (Fig. 3C). *A. fumigatus* did not trigger the Toll pathway within the timeframe upon natural infection, where if anything, *Drs* levels were depressed (Fig. 3D). The low Toll pathway activation by *A. fumigatus* is likely due to the weak ability of this fungus to penetrate insect by the natural route. Collectively, the level of induction of the Toll pathway correlates with the level of pathogenicity using the natural infection mode with *M. anisopliae*>*B. bassiana*>*M. rileyi* >*A. fumigatus*. In sharp contrast, *E. muscae*, despite being very pathogenic to flies, triggers a low induction of *Drs* expression and this upregulation of *Drs* was observed only at the 24 h after infection (Fig. 3E). Expression of another Toll pathway read-out, the Bomanin *BomBc3*, exhibited similar expression to that of *Drs* (Figs. S4A-E). Natural infection of flies carrying the *Drosomycin-GFP* (*Drs-GFP*) reporter (Ferrandon, 1998) also confirmed that *B. bassiana* induced a much stronger expression of the Toll pathway compared to *E. muscae* (Fig. 3G). Thus, natural infection with *E. muscae* either fails to activate the Toll pathway or suppresses it.

**Figure 3.**
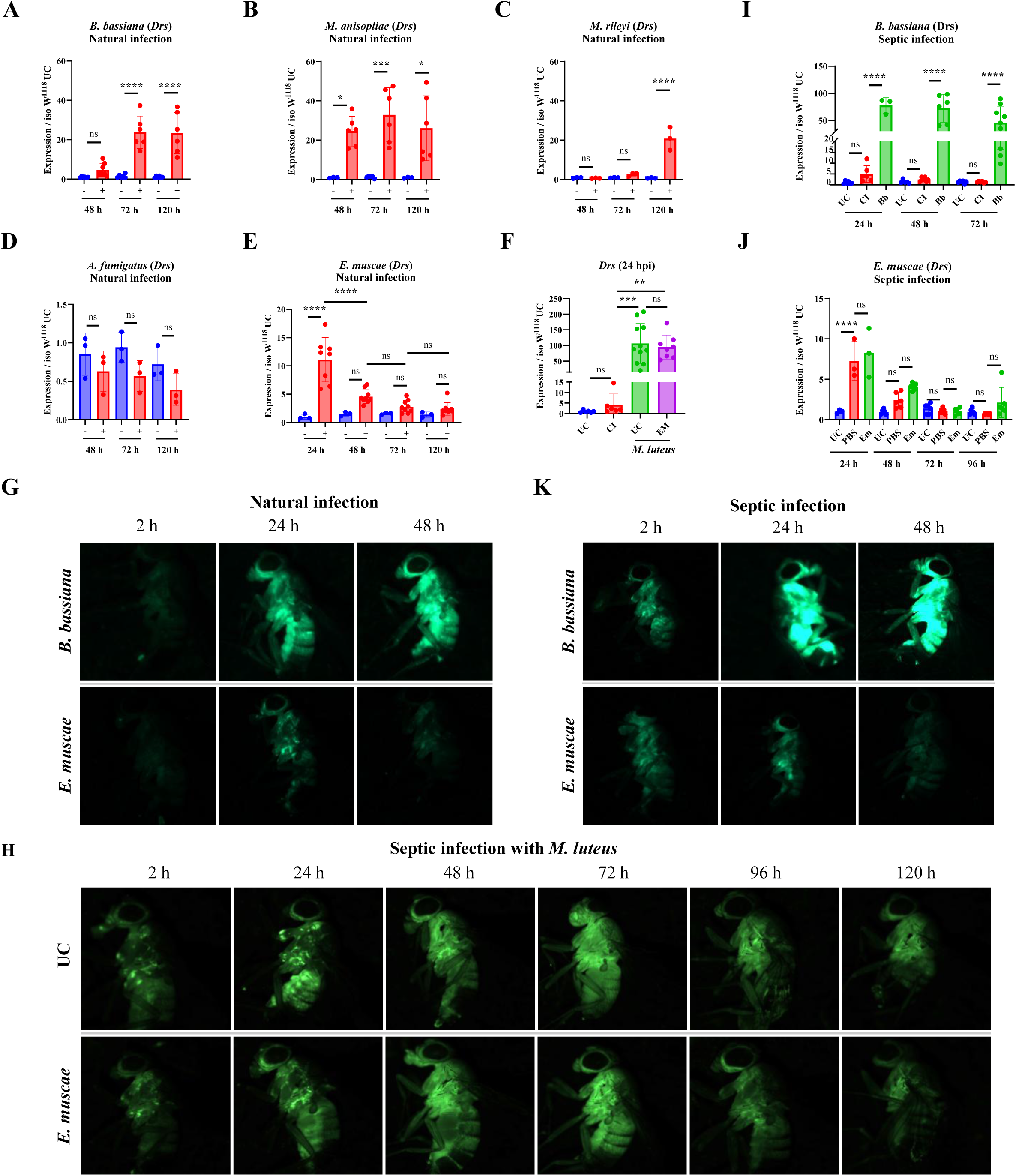
The induction of the Toll pathway in wild-type flies upon infection with the five fungi. (**A-E**) Expression of *Drosomycin* (*Drs*) in wild-type flies upon natural infection with the five fungi. Shown are the relative expression levels of *Drs* in relation to Rp49. *B. bassiana*, *M. anisopliae* and *M. rileyi* induced relatively high *Drs* expression (compare Y axis to F). *A. fumigatus* did not induce *Drs* expression, while *E. muscae* induced a low *Drs* expression level. (**F**) A high induction of *Drs* was observed in unchallenged flies and flies 48 h after natural infection with *E. muscae* upon septic injury with the Gram-positive bacterium *M. luteus*. UC = Unchallenged, CI = Clean Injury, EM = naturally-infected with *E. muscae;* (**G**) Stronger GFP signal intensity was observed in flies carrying the *Drs-GFP* reporter gene upon natural infection with *B. bassiana* than that with *E. muscae*. (**H**) Kinetic analysis of Toll pathway activity was monitored using unchallenged *Drs-GFP* flies and flies 48 h after natural infection with *E. muscae* upon septic injury with the Gram-positive bacterium *M. luteus.* (**I**) Septic injury with *B. bassiana* spores induced higher *Drs* expression than natural infection (Compare I to A). UC = Unchallenged, CI = Clean injection, Bb = *B. bassiana* infection (**J**) Injection with *E. muscae* protoplasts did not induce *Drs* expression (Compare J to E). UC = Unchallenged, PBS = Sterile injection, Em = *E. muscae* injection. (**K**) Stronger GFP intensity was observed in flies carrying the *Drs-GFP* reporter gene upon septic infection with *B. bassiana* than that with *E. muscae*. Expression was normalized with *w^1118^* UC set as a value of 1. Data were analyzed using One-Way ANOVA followed by Tukey’s multiple comparison tests. Values represent the mean ± s.d. of at least three independent experiments. ns, P>0.05, *, P<0.05; **, P<0.01; ***, P<0.001. Full statistical details are available on Table S3.

To distinguish between these two alternatives, we compared the Toll pathway activity of unchallenged flies and flies 48 h after natural infection with *E. muscae* upon septic injury with the Gram-positive bacterium *M. luteus*, a strong inducer of the Toll pathway. We found that 24 h post-infection with *M. luteus, Drs* expression was similarly induced to a high level in both unchallenged flies and flies exposed to *E. muscae* natural infection 48 h prior to *M. luteus* infection, suggesting there was no ongoing suppression of Toll signaling by *E. muscae* (Fig. 3F). Use of flies carrying the *Drs-GFP* reporter revealed similar kinetics of *Drs* expression in 48 h *E. muscae* infected flies and unchallenged flies following infection with *M. luteus* (Fig. 3H). These experiments suggest that natural infection with *E. muscae* fails to activate the Toll pathway rather than suppresses it.

We then compared Toll pathway activity upon septic infection by a needle dipped in spores of *B. bassiana,* or injection of protoplasts of *E. muscae*. As expected, the direct introduction of *B. bassiana* spores triggers a strong *Drs* expression with earlier kinetics compared to natural infection (Fig. 3I). In contrast, injection of *E. muscae* protoplasts did not trigger the Toll pathway more than injury alone as revealed by the low and similar levels of *Drs* expression in both PBS clean injury controls and protoplast-injected flies (Fig. 3J). Expression of another Toll pathway read-out the Bomanin *BomBc3* exhibited similar expression trend to that of *Drs* (Figs. S4F-G). Use of flies carrying the *Drs-GFP* reporter after systemic infection with spores of *B. bassiana* and protoplasts of *E. muscae* further confirmed these observations made by qRT-PCR (Fig. 3K). We infer that protoplasts are not detected by Toll pathway sensors and that this may provide a mechanism for *E. muscae* to proliferate in the hemolymph without triggering the immune response.

Collectively, our study shows that natural infection with *B. bassiana*, *M. anisopliae* and *M. rileyi* triggers a strong induction of the Toll pathway, consistent with their pathogenicity. Despite its pathogenicity, natural infection with *E. muscae* triggers only a low Toll pathway activity indicating that this fungus is not recognized by the immune system.

### *Entomophthora muscae* hides from the host immune response using a vegetative development strategy

To explore the strategies employed by *E. muscae* to hide from Toll sensing, we investigated the dynamics of *E. muscae* development inside flies. We first inspected the steps leading to the protoplast-hypha transition *in vitro* by analyzing protoplast differentiation in Grace’s insect media. We observed that *E. muscae* protoplasts change from a round to an irregular shape (Fig. 4A). The protoplasts grow and elongate towards both ends and other directions along the sites of vesicle of protoplast. Calcofluor White M2R staining showed the absence of cell walls of these protoplasts before their transformation to hyphae after more than two-weeks culture in the Grace’s Insect Medium. We observed the presence of vesicles in both protoplasts and hyphae during the development process in Grace’s insect media (red arrow in Figure 4). Vesicle-driven outward expansion of the membrane ultimately results in pinching of the fungal cell to generate “subprotoplasts”. Vesicles have been demonstrated to promote infection and stimulate protective immunity, depending on the species and host context (Lai et al., 2023; Reis and Rodrigues, 2025; Rodrigues et al., 2024). In addition, DAPI staining reveals that these protoplasts are multinucleated, and in the process of elongating, nuclei disperse along the extension of the protoplasts (Fig. S5). Next, we analyzed protoplast differentiation in flies. Morphology observation of flies showed that the entire infectious process of *E. muscae* could be divided into five stages, namely, early infection stage, fungal spores contact and subsequently penetrate the host cuticle (Stage 1/S1), intra-hemolymph growth stage, protoplasts proliferate inside the flies exhibiting erect wings (S2), protoplasts transformed to invasive hyphae stage, flies affixing themselves with their proboscides just prior to death (S3), hyphal growth stage, dead flies with a swollen and white abdomen with full of hyphae in the hemocoel (S4), hyphae extrude and produce spores from the cadavers (S5) (Fig. 4B). Staining fungal cell by dissecting flies at the above stages S1-S5 with Calcofluor White M2R, which marks fungal chitin in the cell wall, reveals that protoplasts undergo differentiation to hyphae only prior to death in the later stages, with no cell wall at S1 and S2, much thinner cell walls at S3 compared to invasive hyphae at S4 and hyphae sporulating on the cadavers at S5 (Fig. 4C). Microscopic observation showed that the development of *E. muscae* protoplasts also underwent the same morphological steps in host hemocoel as observed *in vitro* (Fig. 4A to Fig. 4C). Although protoplasts undergo the same differentiation process in infected flies, the transition from protoplasts to hyphae takes place 4 to 7 days after fly infection, compared to two weeks in the Grace’s Insect Medium. Thus, the formation of invasive hyphae is a process that take place very late in the infection process, just before fly death. The observation that injection of *E. muscae* protoplasts are not recognized by the immune system, and that *E. muscae* persists in the first stage of infection as protoplasts, suggests that this fungus hides from the immune system by using a cell wall-less structure.

**Figure 4.**
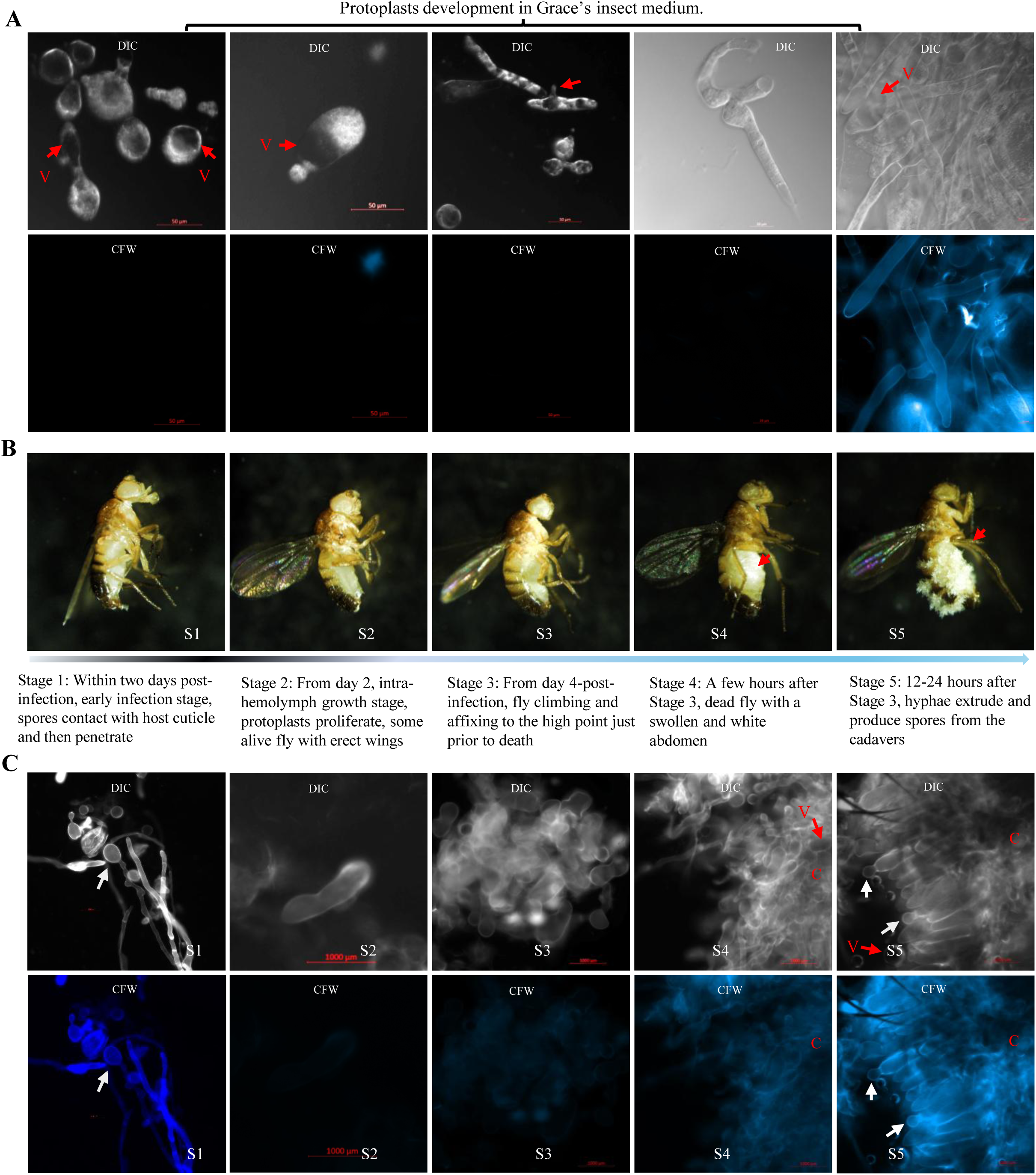
Morphological development of *E. muscae in vitro* culture and in flies. (**A**) The morphological development of *E. muscae* protoplasts in Grace’s Insect Medium showed that round protoplast sprouted and extended to grow at the site of vesicles to form hyphae. Calcofluor White M2R staining showed that cell wall is only present when protoplasts have transformed to hyphae. V: vesicle. (**B**) Five infection stages of *E. muscae* were defined based on the morphological change of flies after infection. Red arrows indicate the white abdomen. (**C**) Calcofluor White M2R staining showed the presence of cell wall of spores contacting with host cuticle (S1), the absence of cell wall of protoplasts (S2), and the presence of thin cell wall when protoplasts transformed to invasive hyphae at S3-S4 and hyphae S5. Red arrows indicate the presence of vesicle in protoplasts and hyphae. C: cuticle; White arrows indicate the primary conidia produced at the tip of the hyphae that extrude from the cadavers.

### The Persephone branch of the Toll-PO serine protease cascade is more critical than the GNBP3-modSP branch to activate the Toll pathway

The Toll pathway is activated upon the maturation of the Toll ligand Spz by a complex cascade of serine proteases called the Toll-PO serine protease (SP) cascade, which also regulates the maturation of PPOs (Westlake et al., 2024). Lysine-type peptidoglycans from Gram-positive bacteria and β-glucans from fungi are detected by the pattern recognition receptors (PRRs) PGRP-SA (with its adaptor GNBP1) and GNBP3, respectively (Gottar et al., 2006; Michel et al., 2001) This triggers the sequential activation of SPs ModSP, Grass, and Psh/Hayan (Buchon et al., 2009; Dudzic et al., 2019; El Chamy et al., 2008; Kambris et al., 2006). Psh and Hayan mediate the activation of SPE, which cleaves Spz, which then dimerizes and binds to the Toll receptor inducing an intracellular cascade leading to the production of Toll-regulated effectors such as antimicrobial peptides (Jang et al., 2006; Weber et al., 2003). Psh (and likely Hayan) can also be independently cleaved by microbial proteases (El Chamy et al., 2008; Gottar et al., 2006; Issa et al., 2018). Hayan and Psh also regulate downstream SPs that activate PPO1 and PPO2, initiating the melanization response leading to cuticular blackening and microbicidal activities (Fig. 5A) (Dudzic et al., 2019; Nam et al., 2012; Shan et al., 2023; Westlake et al., 2024). Having shown the critical role of the Toll pathway upon natural infection with entomopathogenic fungi, we investigated the mechanisms that activate Toll focusing on *B. bassiana*, *M. anisopliae, M. rileyi* and *E. muscae*. In these experiments, we used flies carrying the *GNBP3^hades^* null mutation over a deficiency *Df(ED4421),* as we noticed that *GNBP3^hades^*homozygous flies display a marked susceptibility to fungal infection that was not observed in transheterozygote *GNBP3^hades^/Df(ED4421)* flies, pointing to a background effect (Figs. S6A and 6B). The marked susceptibility of the *psh^SK1^* analysis showed that Psh contributes to host defense against *B. bassiana*, *M. anisopliae, M. rileyi* and *E. muscae* (Figs. 5B-E). In contrast, the role of the glucan binding sensor GNBP3 was modest as revealed by the almost wild-type survival of *GNBP3^hades^/Df(ED4421)* flies to *B. bassiana*, *M. anisopliae, M. rileyi* and *E. muscae* (Figs. 5B-E, Table S3). Similar results were obtained when using another recently generated *GNPB3^Δ^*deficient mutant (Lu et al., 2024) (Figs. S6A and S6B). As ModSP integrates signals from GNBP3 as a PRR of the Toll-PO SP cascade (Buchon et al., 2009), we expected that *modSP^1^* flies would exhibit the same host defense as *GNBP3^hades^/Df (ED4421)* flies against fungal infection. We indeed observed that *modSP^1^* flies exhibited the same ability as *GNBP3/Df (ED4421)* flies to survive *B. bassiana*, *M. anisopliae* and *M. rileyi (*Fig 5B-D). Strikingly, *modSP^1^*flies but not *GNBP3^hades^/Df(ED4421)* displayed a marked susceptibility to *E. muscae* similar to *spz^rm7^* flies (Fig 5E). As expected, *psh^1^;;modSP^1^* double mutants that block both modes of sensing of Toll and *Hayan-psh^Def^* double mutants displayed susceptibility similar to *spz^rm7^* flies. To further characterize the role of the Toll sensor, we next monitored Toll pathway activation upon natural infection with *M. anisopliae*. We restricted our analysis to this fungal species as *M. anisopliae* strongly activates the Toll pathway. The expression pattern of *Drs* and *BomBc3* in the mutants confirms the main role of Psh in activation of the Toll pathway upon natural infection (Figs. 5F-G). Both *Drs* and *BomBc3* expression were not induced in *psh^SK1^* flies, *Hayan*-*psh^Def^* double mutant flies, or *psh^1^;;modSP^1^* double mutant flies, being comparable to *spz^rm7^* flies, consistent with their high susceptibilities to fungal infection. However, the expression of the Toll pathway was not affected in *GNBP3^hades^* homozygous flies as revealed by the wild-type expression of *Drs* and *BomBc3*.

**Figure 5.**
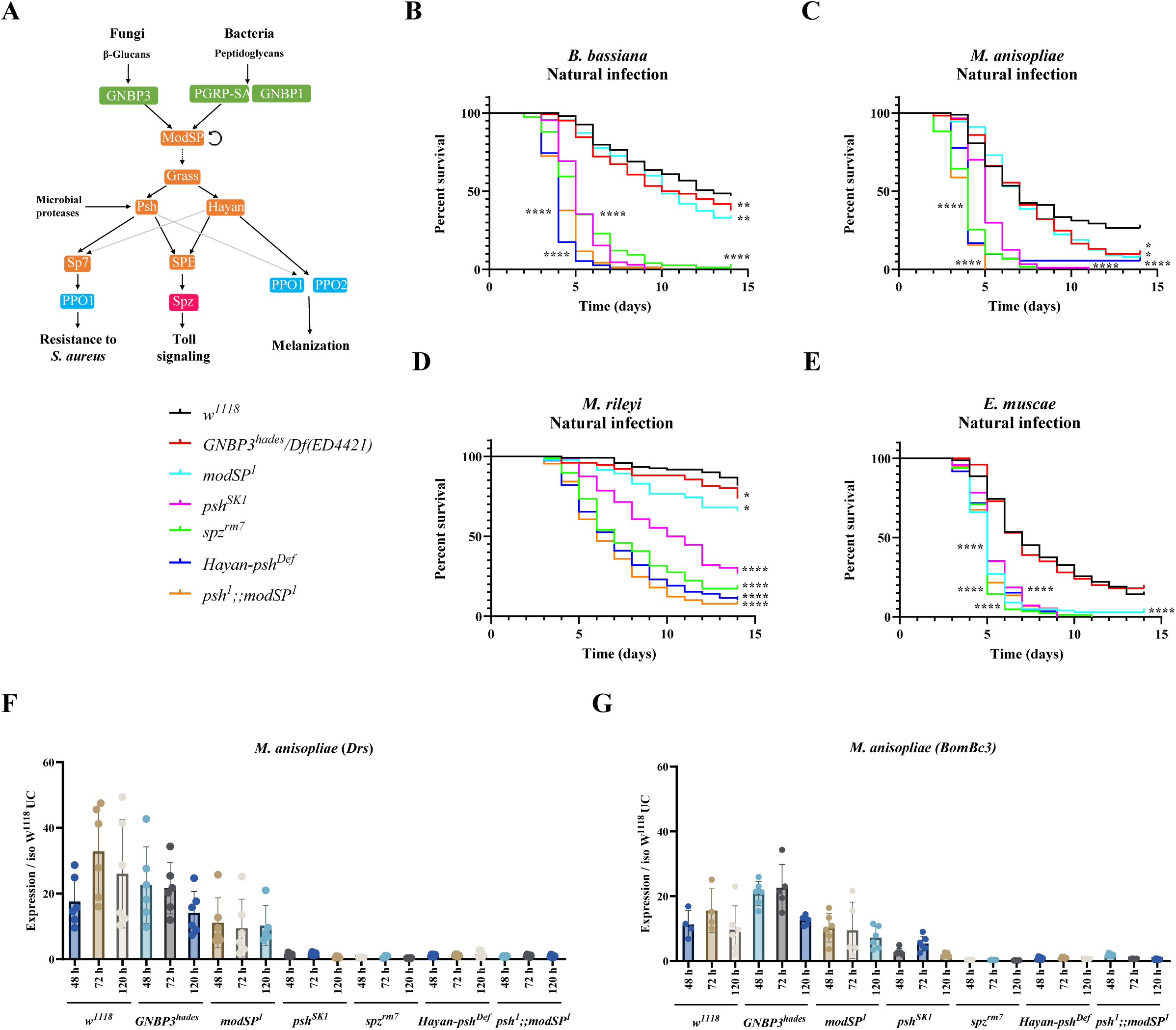
Contribution of the Psh and GNBP3 to Toll pathway activation. (**A**) A model of GNBP3 and SPs regulating the Toll pathway and the Melanization reaction upon fungal infections. (**B-E**) Natural infection with *B. bassiana*, *M. anisopliae*, and *M. rileyi* revealed that *psh^SK1^* flies have a reduced survival rate compared to *GNBP3^hades^/Df (ED4421)*, *modSP^1^* and wild-type flies (P<0.0001). *modSP^1^* flies were as susceptible to *E. muscae* as *spz^rm7^* and *psh^1^* flies (P>0.05). The Hayan-*psh^Def^* double mutant flies is more susceptible to *B. bassiana*, *M. anisopliae* and *M. rileyi* than *psh^SK1^* (Bb, χ²=36.19, P<0.0001; Ma, χ²=16.91, P<0.0001; Mr, χ²=14.14, P=0.0002). Likewise, the double mutant flies *psh;;modSP* were more susceptible to *B. bassiana*, *M. anisopliae* and *M. rileyi* than *psh^SK1^* (Bb, χ²=14.65, P=0.0001; Ma, χ²=54.88, P<0.0001; Mr, χ²=22.68, P<0.0001) and *modSP^1^* (Bb, χ²=173.1, P<0.0001; Ma, χ²=120.6, P<0.0001; Mr, χ²=55.81, P<0.0001). (**F**) Expression of *Drs* revealed that *GNBP3^hades^* flies have wild-type *Drs* expression levels (p>0.05). The Toll pathway was not activated in *psh^SK1^* flies as in *spz^rm7^*, *psh^1^* and *Hayan-psh ^Def^* flies, and displayed a lower induction in *modSP^1^* flies compared to wild-type flies. (**G**) Expression of *BomBc3* showed similar results to the expression of *Drs*. The *spz^rm7^* flies as a positive control. Survival tests were analyzed by Log-rank test and gene expression data were analyzed using One-Way ANOVA followed by Tukey’s multiples comparison tests. Values represent the mean ± s.d. of at least three independent experiments. Full statistical details are available on Table S3.

Our results indicate that Psh is a highly important sensor in activating the Toll pathway to mediate the host defense against these fungi, while GNBP3 plays only a minor role. Surprisingly, we note that ModSP plays a specific role in surviving infection with *E. muscae, suggesting another role of this apical serine protease beyond integrating GNBP3 signals*.

### The Toll pathway restricts fungal growth

Having characterized the importance of the Toll pathway and the underlying mechanism leading to its activation, we next explored how the Toll pathway effectively combats fungal infection. Protection by the immune system can be mediated via resistance mechanisms that prevent pathogen growth or via tolerance mechanisms that protect the host without directly inhibiting the pathogen (Pradeu et al., 2024; Soares et al., 2017). We infected flies by natural infection and quantified fungal proliferation after infection. qPCR levels of *B. bassiana 18S* gDNA reveal a much high fungal load at 3- and 4-days post natural infection in *spz^rm7^* flies compared to wild-type flies (Fig. 6A). Higher fungal loads with earlier growth kinetics were also found in *spz^rm7^* flies compared to wild-type flies when infected with *M. anisopliae* and *E. muscae* (Figs. 6B-C). To confirm and extend these results, we used a *M. anisopliae* tagged GFP strain and monitored the expression of the Green Fluorescent Protein (GFP) upon septic infection in both wild-type and *spz* mutant flies. We observed a much higher proportion of flies with strong GFP signals in *spz^rm7^* flies compared to wild-type (Fig. 6D). While *M. anisopliae* disseminated throughout the body in *spz^rm7^* mutant flies, the fungus was restricted to the thorax in wild-type flies at 2- and 3- days post infection (Fig. 6E). In keeping with this faster dissemination, the fungi broke through cuticle and extrude from the cadavers of *spz^rm7^* mutant flies 2 to 4-day before that of *wild-type* flies. In both wild-type and *spz^rm7^* flies, fungal growth eventually occurs in the cadavers across all three tagmata, including the wings and legs (Fig. S7). Thus, Toll act as a resistance mechanism limiting the growth of these three entomopathogenic fungi.

**Figure 6.**
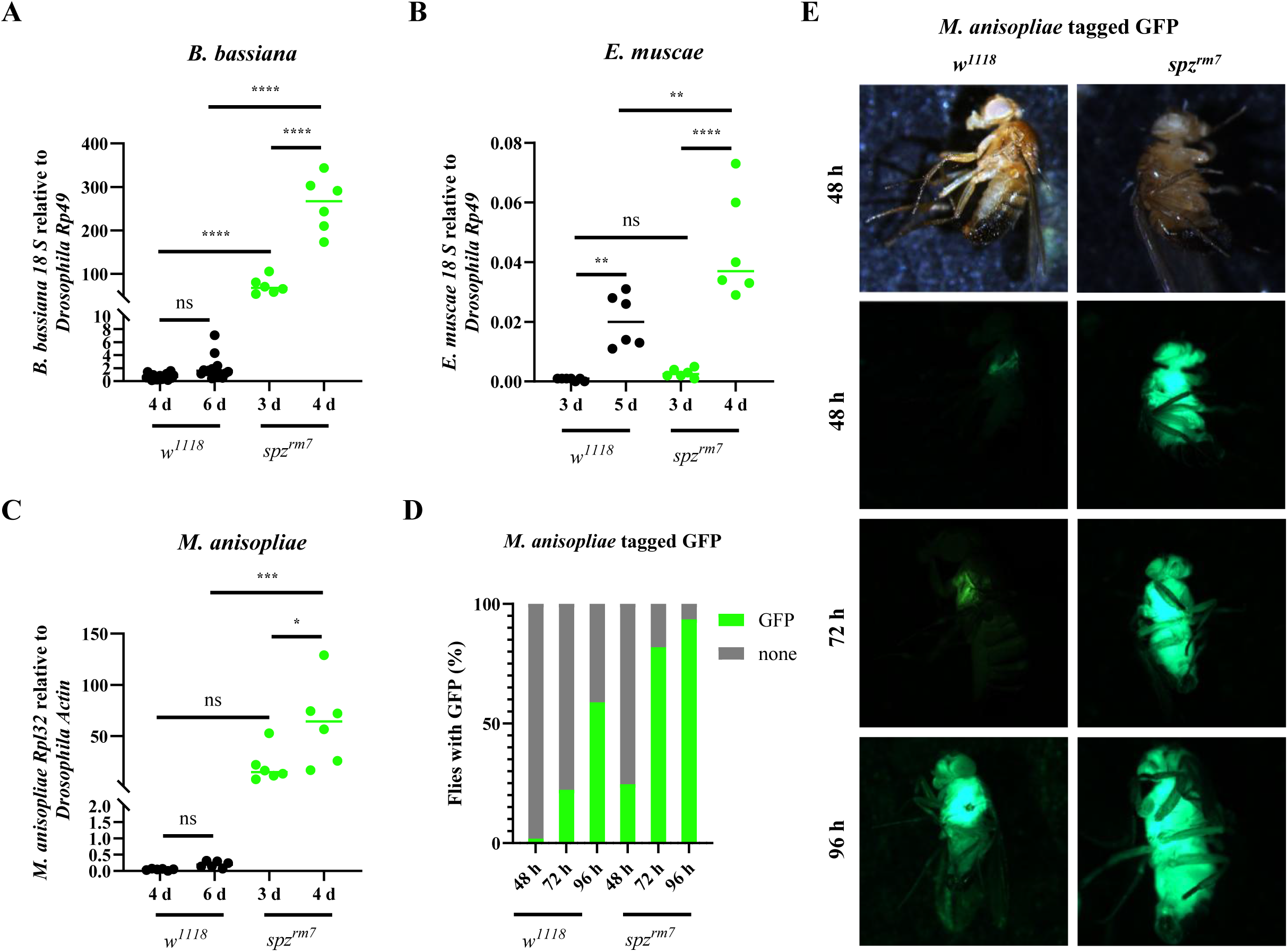
Higher fungal persistence in flies deficient for the Toll pathway. (**A-C**) Quantification of fungal loads by qPCR analyses reveals higher fungal loads in *spz^rm7^* flies compared to wild-type flies after natural infection with *B. bassiana*, *M. anisopliae* and *E. muscae*. (**D-E**) Higher proportion of flies with GFP and a stronger GFP intensity were observed in *spz^rm7^* flies compared to wild-type flies at 48, 72 and 96 hours after pricking with spores of a *M. anisopliae* tagged GFP strain. ns, P>0.05, **, P<0.01; ****, P<0.0001. Full statistical details are available on Table S3.

### Multiple host defense peptides contribute to Toll antifungal activity

In the last years, several host new defense peptides controlling fungal infection have been identified in *Drosophila* (Hanson et al., 2025). In addition to Drs and Mtk that display antifungal activity *in vitro* (Feldman et al., 2019; Moghaddam et al., 2017), genetic studies have revealed that deficiencies in *Baramicin A*, *Bomanins* (12 genes) and *Daisho* (2 genes) cause susceptibility to specific fungi (Clemmons et al., 2015; Cohen et al., 2020; Hanson et al., 2021). The role of these immune peptides has never been analyzed collectively for their role in host defense against fungi. This prompted us to explore the nature of downstream Toll effectors using isogenic mutant lines against entomopathogenic fungi upon natural infection. In this analysis we used single null mutants in *Drosomycin (Drs^R1^)*, *Metchnikowin (Mtk^R1^)*, *Baramicin A (BaraA^SW1^)*, *Daisho1/2* (*△Dso*, a short deletion removing both Daisho 1 and 2) and *Bom^△55C^* that removes ten of the twelve *Bomanins* (Clemmons et al., 2015; Cohen et al., 2020; Hanson et al., 2021, 2019). We also used previously generated compound mutants including *Mtk ^R1^, Drs^R1^* double mutants, *△*AMP10 (a fly line deleted for 10 AMPs including *Mtk*, *AttA/B/C/D*, *DptA/B*, *Drc*, *Def*) and *△*AMP14 (a fly line deleted for 14AMPs similar to *△*AMP10 with an additional deletion of the four *Cecropin* genes *CecA1/A2/B/C*) flies (Carboni et al., 2022; Hanson et al., 2019). In the framework of this study, we further generated four additional compound mutants to see possible additivity or synergistic action of these host defense peptides including i) *BaraA ^SW1^*, *△Dso*; ii) *Mtk^R1^*, *BaraA^SW1^*; *Drs^R1^*; iii) *Mtk ^R1^, BaraA^SW1^*, *△Dso*; *Drs^R1^*) and iv) *Mtk ^R1^*, *Bom^△55C^*; *Drs^R1^*.

Upon natural infection by fungi, the *Bom^△55C^*, *△Dso* deletions caused marked susceptibility to all four entomopathogenic fungi while single mutations in *Mtk* and *Drs* had no effect against any fungus. Alone, *BaraA^SW1^* caused a susceptibility to only two of the four fungi. This reveals the critical role of *Bomanin* and *Daisho1/2* to survive fungal infection (Fig. 7A and Fig. S8). We found that flies lacking both *Baramicin* and *Daisho1,2* were markedly more susceptible to infection with *B. bassiana, M. rileyi,* and *E. muscae*, compared to *△Dso* single mutant flies, while the difference for *M. anisopliae* was consistent in direction, but relatively minor (Table S3). We also observed that *Mtk^R1^*, *Drs^R1^* but not *Mtk^R1^*or *Drs^R1^* single mutants display increased susceptibility to *M. anisopliae,* but not *B. bassiana*, *M. rileyi* and *E. muscae, revealing an additive effect against M. anisopliae specifically*. Strikingly, *Mtk ^R1^*, *Bom^△55C^*; *Drs^R1^* showed greater susceptibility to infection compared to *Bom^△55C^* flies for *B. bassiana, M. anisopliae,* and *M. rileyi,* but not *E. muscae*. The susceptibility of *Mtk^R1^*, *Bom^△55C^*; *Drs^R1^* flies was slightly but consistently less marked than *spz^rm7^* mutant flies across all fungi. Finally, we observed that *△AMP14* flies display greater susceptibility compared to *△AMP10* flies against each fungus, pointing to a role of Cecropin in host defense against fungi, consistent with *in vitro* observation (Ekengren and Hultmark, 1999).

**Figure 7.**
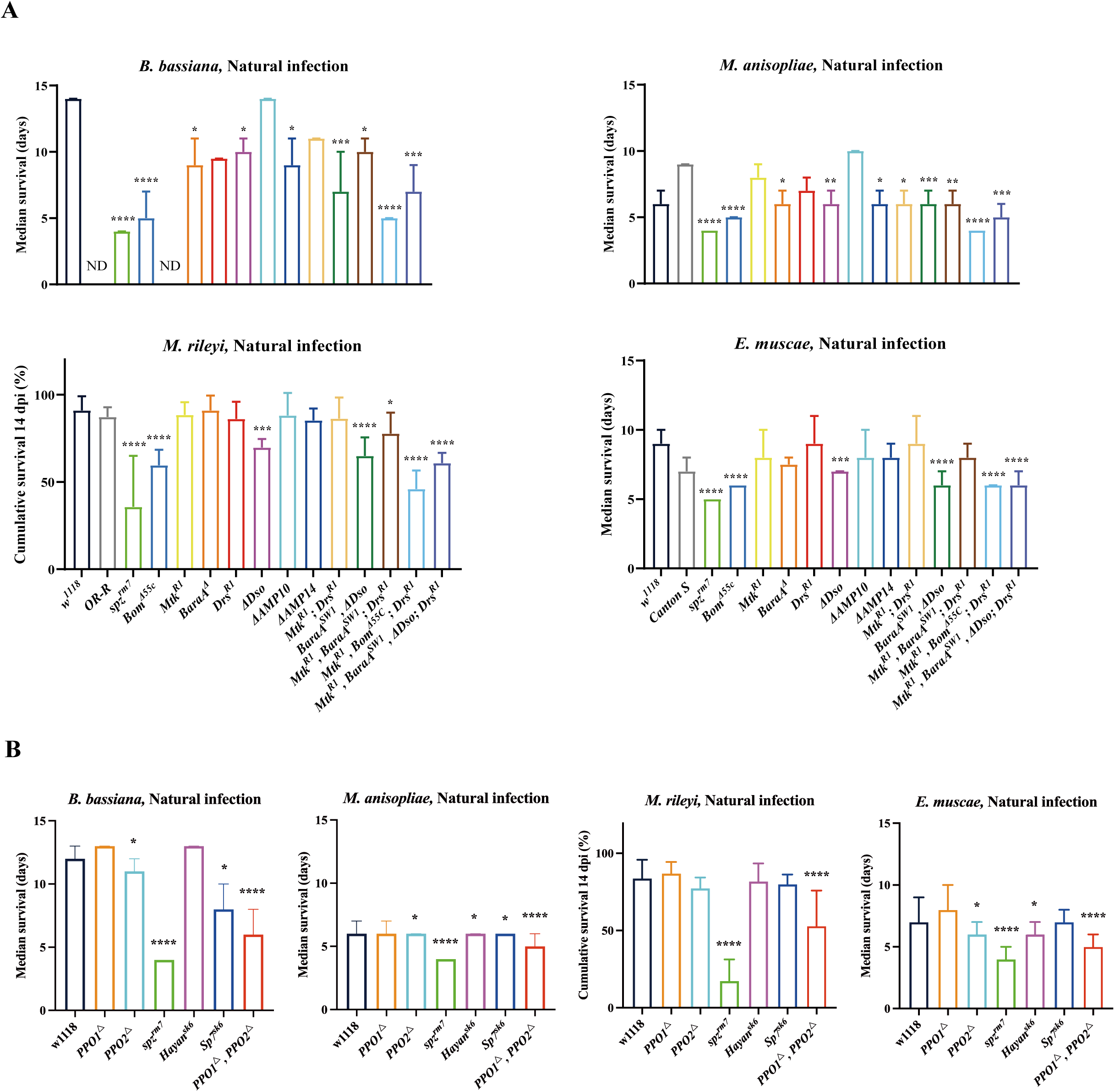
Contribution of host defense peptides and Serine Proteases involved in melanization to survival upon fungal infection. (**A**) The Bomanin and Daisho1/2 are critical for defense against *B. bassiana*, *M. anisopliae*, *M. rileyi* and *E. muscae* infection. (**B**) PPO2 mutant flies were more susceptible to fungal infection than PPO1. The *spz^rm7^* flies as a positive control. Shown are the median survival for infection with *B. bassiana*, *M. anisopliae* and *E. muscae*, and cumulative survival for infection with *M. rileyi*. Data were analyzed using the Log rank test. ns, P>0.05; *, p<0.05; ***, P<0.001; ****, P<0.0001. Full statistical details are available on Table S3.

Taken together, we show that host defense peptides regulated by the Toll pathways, notably Bomanins, Daisho1/2, BaraA, Mtk and Drs largely explain the contribution of Toll to survive fungus natural infections. The strength of effect was broadly consistent across fungal species, with virulent species revealing a greater resolution of differential susceptibilities. Generally, we did not find evidence for a specific importance of any fungus-by-gene interactions among these effectors, suggesting their contributions to survival are based on their collective action in antifungal defense, with some being more important than others.

### Both PPO1 and PPO2 additively contribute to host defense against fungi

Having described the contribution of Toll effectors, we next analyzed how melanization contributes to host defense against these entomopathogenic fungi. Previous studies have shown that PPO1 and PPO2 redundantly contribute to both cuticular and hemolymphatic melanization (Binggeli et al., 2014; Dudzic et al., 2019). Genetic approaches have allowed us to distinguish distinct activities associated with PPO1 and PPO2: cuticular melanization at the injury site is dependent on PPO1 activated through Hayan, while an uncharacterized microbicidal activity against *Staphylococcus aureus* relies on Sp7, but not Hayan, with a major role for PPO1 and a lesser but important role for PPO2 (Dudzic et al., 2019); we will note, however, that there is some redundancy *in vitro* regarding the capability of these SPs to cleave PPO1 and PPO2 (Shan et al., 2023). Surprisingly, compared to these roles against *S. aureus,* we confirmed a consistent but opposite trend in the importance of *PPO1* and *PPO2* in defense against natural infection by each fungus (Fig. 7B and Fig. S9). *PPO2* mutants were more susceptible to all fungi compare to PPO1, although *PPO1, PPO2* double mutants were consistently more susceptible than flies deleted for *PPO2* alone (Table S3). We next tested deletions in *Hayan* (*Hayan^SK6^)* and *Sp7* (*Sp7^SK6^*). Agreeing with a primary role of Sp7 in PPO2-mediated defense, *Sp7^SK6^* flies were as susceptible to natural infection by *B. bassiana* and *M. anisopliae* as *PPO2* mutants. On the other hand, *Hayan* mutation caused no susceptibility to *B. bassiana,* but a similar susceptibility to *Sp7* and *PPO2* mutants to *M. anisopliae.* No single mutation caused a significant mortality against *M. rileyi.* Strikingly, *Hayan* mutation, but not *Sp7*, had a comparable susceptibility to *PPO2* mutants to *E. muscae.* In all cases, *spz^rm7^* was the most susceptible genotype.

We also monitored fungal load of *B. bassiana* and *M. anisopliae* in these genotypes at key time points after infection. qPCR of *B. bassiana 18S* and *M. anisopliae Rpl32* revealed trends broadly consistent with survival curves, with higher fungal loads broadly correlating to survival (Figs. 8A-B). We also infected wild-type and melanization deficient flies by pricking with *M. anisopliae* GFP spores in the thorax. In these experiments bypassing the cuticle, we again observed faster growth of *M. anisopliae GFP* in *PPO1^△^*, *PPO2^△^* flies revealed by stronger GFP signals and higher dissemination level within the body at both 2- and 3-day post-infection compared to wild-type flies (Fig. 8C). Consistent with the survival analysis, we observed slightly faster growth *M. anisopliae-GFP* in *PPO2* single mutants compared to *PPO1* single mutants, with a slightly earlier and stronger GFP signal also seen for *Hayan* and *Sp7* deficient flies both at 48 h and 72 h after pricking compared to wild-type and *PPO1* mutants (Fig. 8C).

**Figure 8.**
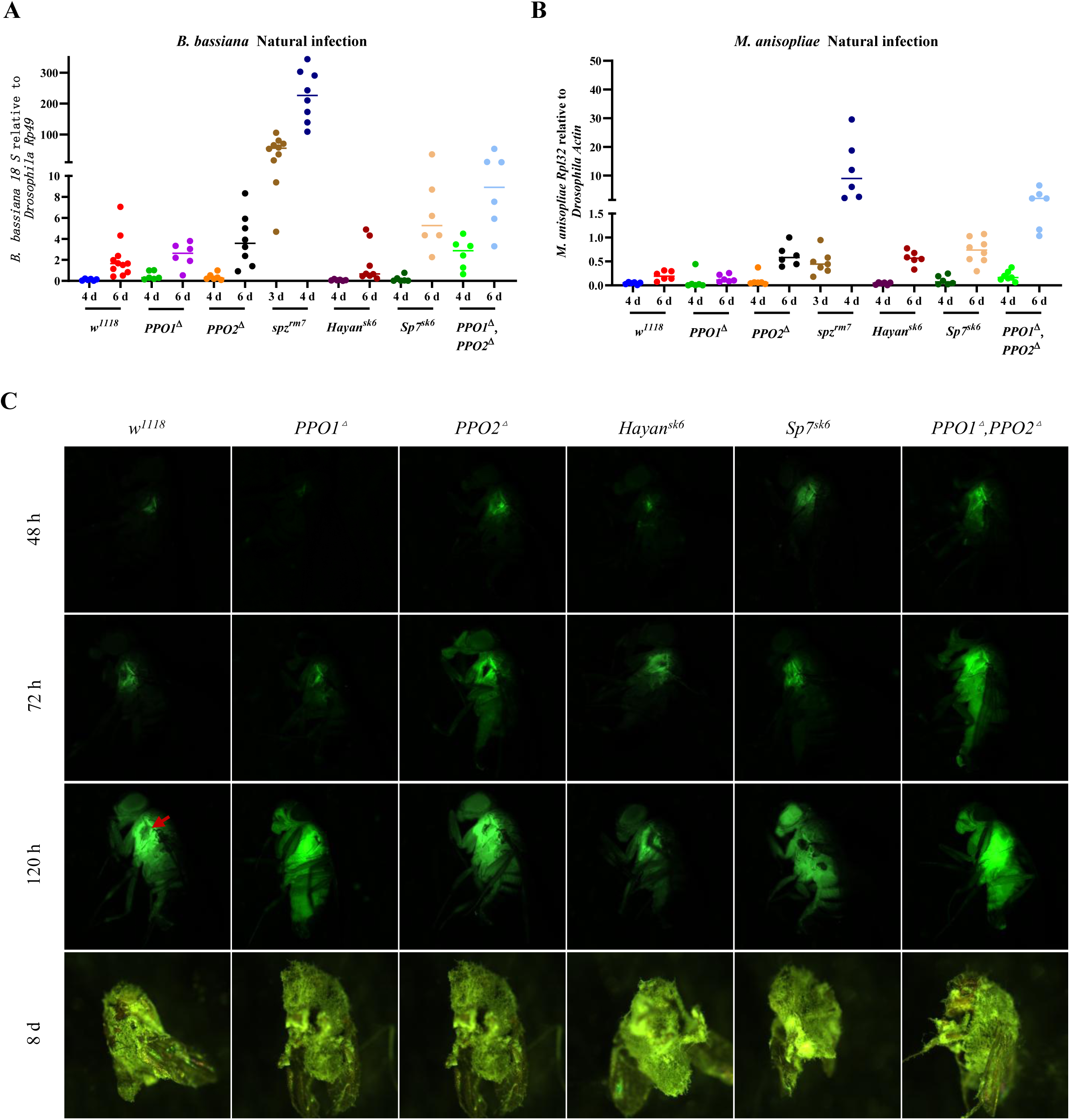
Melanization limits fungal growth. (**A**) *B. bassiana* fungal load increases in wild-type and mutants before mortality upon natural infection. *spz^rm7^* and *PPO1^△^, PPO2 ^△^* flies have a higher fungal persistence in *PPO1 ^△^, PPO2 ^△^* flies. (**B**) *M. anisopliae* fungal load increases in wild-type and mutants showing similar trends as *B. bassiana* fungal load upon natural infection. Data were analyzed using One-Way ANOVA followed by Tukey’s multiples comparison tests and values are pooled data from three independent experiments. (**C**) Stronger GFP intensity and faster fungal dissemination was observed in *spz^rm7^* and *PPO1^△^, PPO2^△^* flies upon pricking with spores of a *M. anisopliae* tagged GFP strain. Red arrow indicates the growth of hyphae at the melanization spot of the pricking site. Full statistical details are available on Table S3.

Collectively, we observe a stronger role of PPO2 compared to PPO1 in defense against fungi. Additionally, it reveals different requirements for Sp7 and Hayan in defense against these fungal species, somewhat agreeing with previous observations for the role of these serine proteases in activating PPO1 or PPO2 in defense against *S. aureus,* although there is some additional complexity to the antifungal response. There was a strong correlation between susceptibility and fungal growth in the tested mutants, indicating that melanization contributes to host survival by limiting fungal dissemination and growth.

### Higher Toll pathway activity in *PPO1^△^*, *PPO2^△^*mutant

Finally, we explored whether the immune susceptibility of melanization defective flies might result from improper Toll activity in the absence of melanization, we monitored the activation of the Toll pathway in *PPO1*, *PPO2* single and double mutant flies as well as *Sp7* and *Hayan* flies. Single mutations in *PPO1*, *PPO2*, *Sp7* and *Hayan* did not affect the inducibility of *Drosomycin* upon natural infection with *M. anisopliae* and *B. bassiana* (Figs. S10A-B). In agreement with previous studies (Binggeli et al., 2014; Ryckebusch et al., 2025), we found a trend of higher Toll pathway activation in *PPO1^△^*, *PPO2^△^* double mutant flies compared to *wild-type* flies upon natural infection with these two fungal species, although *PPO* single mutants showed no increased Toll activity. A higher and earlier Toll pathway activation was also observed upon direct introduce of *M. anisopliae* spores in the body cavity at the time points of 24 h and 48 h, but not 72 h (Fig. S10C). Interestingly, we also observed a higher level of *Drs* expression in *PPO1^△^*, *PPO2^△^* flies compared to wild-type flies upon injury with a clean needle (Fig. S9C). Moreover, unchallenged *PPO1^△^, PPO2^△^* flies tend to have higher basal expression of *Drs* compare to their wild-type counterpart across each time point (Fig. S10C).

These findings indicate that higher Toll signaling in *PPO1^△^, PPO2^△^* is not strictly linked to higher fungal load, but may relate to a negative feedback loop between the Toll and the melanization cascade. This negative feedback loop has been proposed previously using live and heat-killed microbe injections (Binggeli et al., 2014; Ryckebusch et al., 2025). Here we show this increased Toll activity is true even for unchallenged and clean injury treatments, suggesting a role for PPOs in regulating Toll activation even in the absence of infection.

## Discussion

Host–fungal pathogen interactions are complex and serve as potent drivers of evolutionary diversification (Brown, 2023; Gladieux et al., 2014; Murante and Hogan, 2024; Shang et al., 2016). Fungal pathogens span a broad ecological spectrum-from generalists like *B. bassiana* and *M. anisopliae* with wide host ranges, to specialists such as *M. rileyi*, *E. muscae*, and *O. sinensis*, to opportunistic fungi like *C. albicans* and *A. fumigatus*. Despite this diversity, our understanding of how fungal pathogens with distinct lifestyles interact with host immunity remains incomplete (Lu et al., 2024; Murante and Hogan, 2024; Qu and Wang, 2018; Wang and Wang, 2017).

To address this gap, we used *D. melanogaster* fly lines deficient in specific immune pathways and effectors (Hanson et al., 2023, 2019; Ryckebusch et al., 2025) to systematically investigate host defense against five fungal species. Consistent with earlier work (Lemaitre et al., 1996; Ryckebusch et al., 2025; Westlake et al., 2024), the Toll pathway emerged as the most critical immune module for defense against all tested fungi. Generalist pathogens like *B. bassiana* and *M. anisopliae*, and to a lesser extent the specialist *M. rileyi*, strongly induced the Toll pathway reporter gene *Drs* upon natural infection. In contrast, *E. muscae* triggered only weak activation, and *A. fumigatus* did not induce *Drs* expression at all upon natural infection. Nevertheless, Toll signaling remained essential for survival even against weak inducers, suggesting that basal or low-level Toll activity is sufficient to confer protection.

Our findings further show that Toll pathway-mediated immunity is important not only after natural infection, which involves cuticle penetration, but also after systemic infections when fungal spores are introduced directly into the hemolymph. This emphasizes the role of the Toll pathway in limiting fungal growth systemically. Recent studies using deletions of immune effectors downstream of the Toll and Imd pathways have demonstrated synergistic, additive, and pathogen-specific roles in survival (Hanson et al., 2025, 2019). We show here that Bomanins and Daisho peptides are essential for surviving infection by generalist and specialist entomopathogenic fungi. These observations align with previous studies showing the broad antifungal activity of Bomanins against *Fusarium spp.*, *B. bassiana*, *M. anisopliae*, and yeast species (Clemmons et al., 2015; Hanson et al., 2019), and Daisho peptides active against *Fusarium* species and some *Aspergillus* strains (Cohen et al., 2020). In addition, Baramicin A peptides are important for defense against virulent *Beauveria* and *Metarhizium* species (Hanson et al., 2021; Huang et al., 2023), while an individual importance for Drosomycin and Metchnikowin was not observed for any fungus tested (Hanson et al., 2021, 2019). However, the Baramicin, Drosomycin, and Metchnikowin peptides each contribute to defense independently from other Toll effectors, evidenced by an increased mortality of *BaraA, Dso* and *Drs, Mtk, Bom* combinatory mutants against most fungi. Our results may help make sense of previous studies showing striking single-mutant susceptibilities (Cohen et al., 2020; Hanson et al., 2021), as the importance of these genes was revealed in a virulence-dependent, or host susceptibility-dependent, manner – virulence and susceptibility broadly measuring opposite sides of the same coin. Thus, unlike the specificity observed for antibacterial peptides (Hanson et al., 2023, 2019; Unckless et al., 2016), the antifungal peptides we tested appear to act in a more general fashion, suggesting that they target broadly conserved fungal features. The similar susceptibility profiles of *BaraA^SW1^, Mtk^R1^*, *△Dso; Drs^R1^* mutants to *Bom^Δ55C^* underscores the critical role of these humoral Toll-mediated effectors in antifungal immunity, with additional mutations (e.g. *Mtk^R1^*, *Bom^Δ55C^; Drs^R1^* flies) increasingly approaching *spz^rm7^* levels of susceptibility. It would be interesting to achieve the full Toll effector combinatory mutant line (*BaraA^SW1^, Mtk^R1^, Bom^Δ55C^, △Dso; Drs^R1^*) to learn if there is any host defense mediated by *spz* beyond the contributions of these five effector families.

As our *Bom^Δ55C^* and *△Dso* deletions removed large gene clusters (10 of 12 Bomanins and both Daisho genes), we could not assess individual gene contributions. However, recent work has identified roles for specific Bomanins such as BomT1 and BomT2 in surviving *Enterococcus faecalis*, *M. robertsii*, and *A. fumigatus* ribotoxins (Lou et al., 2025). Identifying the Bomanins that mediate defense against the fungi tested here will be an important next step. In addition, it would be interesting to confirm the precise mechanism through which Cecropins improve survival against entomopathogenic fungi. While these peptides can display antifungal activity *in vitro* (Ekengren, 1999). The protective effect of Cecropins could therefore rely either on a direct antifungal action, or an indirect role in preventing dysbiosis, perhaps also modulating the cuticle microbiota that itself protects against topical fungal infections (Hong et al., 2022).

Unexpectedly, we did not observe a major role of the glucan sensor GNBP3 in activating Toll in response to *M. anisopliae*. A recent study showed that GNBP-like3 (GL3) acts as a bait for the fungal effector Tge1, which can inhibit GNBP3 and so suppress Toll immune activation (Lu et al., 2024); itself does not propagate Toll signaling. However, our results using *ΔGL3*, *ΔGNBP3* or *GNBP3^hades^/Df(ED4421)* flies indicate a minor role only for this GNBP3 arm of Toll signaling in survival upon *B. bassiana* and *M. anisopliae* natural infection (Fig. S6). Instead, our data support a predominant role for the Persephone (Psh) branch in Toll activation, suggesting that entomopathogenic fungi are primarily recognized via the proteases they secrete (Gottar et al., 2006). These include serine proteases and chitinases involved in cuticle degradation (Qu and Wang, 2018), to which Psh may be specifically adapted.

Melanization also contributed to host defense, albeit to a lesser extent than Toll signaling, consistent with earlier findings (Binggeli et al., 2014; Ryckebusch et al., 2025). Its reduced impact upon septic infection suggests a key role at the initial cuticular breach, where melanin and ROS deposition may restrict fungal entry and dissemination. This is consistent with observations in *Galleria mellonella* and *Anopheles gambiae*, where melanin accumulates at fungal entry sites (Vertyporokh et al., 2020; Yassine et al., 2012).

We show that both PPO1 and PPO2 contribute to fungal resistance. Interestingly, across fungal pathogens we found PPO2 plays a more prominent role in host defense compared to PPO1, even after septic injury with *M. anisopliae*. Previous studies have described a major role for PPO1 in wound site melanization through the action of Hayan, but also PPO1 in host defense against *S. aureus* in a Hayan-independent manner, with a more minor role for PPO2 (Binggeli et al., 2014; Dudzic et al., 2019; Vasanth et al., 2025). However, here using entomopathogenic fungal infections, we found the opposite. Unlike the case of *S. aureus,* distinct roles for *Hayan* and *Sp7* in this PPO2-mediated immune defense are less clear, perhaps reflecting an importance of Sp7 in the hemolymph, and of Hayan as the PPO-converting SP at the injury site as suggested by its essentiality for wound blackening (Dudzic et al., 2019); however, wound site blackening requires Hayan acting through PPO1. We also observed higher Toll pathway activation in *PPO1, PPO2* double mutant flies compared to wild-type flies upon fungal natural infection, agreeing with previous studies (Binggeli et al., 2014; Ryckebusch et al., 2025). Here we additionally show this higher Toll activation upon clean injury and even in unchallenged flies. This suggests that byproducts of POs somehow negatively regulate Toll activation, which include terminal reactive oxygen species, but also various intermediates involving the conversion of tyrosine to L-DOPA and derivatives such as dopaquinone or the production of melanin from 5,6-dihydroxyindole (DHI) (Westlake et al., 2024). Conversely, Dopa decarboxylase converts L-DOPA to dopamine, and so the deficiency of PO activity converting L-DOPA to dopaquinone could result in the inverse accumulation of dopamine, likely having pleiotropic effects. Here, the higher Toll activity in *PPO1, PPO2* double mutant flies suggests there is effectively negative feedback exerted by the melanization pathway on the Toll pathway. It would also be interesting to know if this role of melanization in Toll regulation is related to the role of both modules in the wound healing response (Capilla et al., 2017; Carvalho et al., 2014).

The Imd pathway had minimal roles in host defense against the four entomopathogenic fungi tested, except during *A. fumigatus* septic infection. Interestingly, volatile organic compounds from *A. fumigatus* have been shown to be toxic to Imd-deficient flies (Almaliki et al., 2023), and a previous study showed that *A. fumigatus* virulence acted through either proliferation or secretion of toxins in a Bomanin and melanization-dependent manner (Xu et al., 2023), suggesting *A. fumigatus* deploys multiple pathogen strategies that interact with different arms of host immunity.

Previous studies have showed that entomopathogenic fungi have evolved sophisticated strategies to evade phagocytosis by hemocytes. Once fungal spores (conidia) germinate, penetrate host tegument and reach the hemocoel, fungi exist within the hemocoel in the forms of blastospores with thinner cell walls than conidia (*M. anisopliae*, *M. rileyi*, *B. bassiana*), and cell wall-free protoplasts (*E. muscae*). (Wang and St. Leger, 2006) had demonstrated that host hemocytes can ingest conidia of *M. anisopliae*, but are unable to recognize blastospore, due to the production of a cell surface hydrophobic protein gene *Mcl1* that is expressed within 20 min of the fungal pathogen contacting hemolymph. Studies have shown that blastospores of *B. bassiana* and *M. anisopliae* can be phagocytosed at the early stages of infection but manage to escape from host cells and continue to propagate. Growing hyphal bodies can deform the plasmatocyte cell membrane, and the hemocyte count gradually decreases as observed in Lepidopteran insect during fungal infection (Gillespie et al., 2000; Hung and Boucias, 1992; Li et al., 2020a; Liu et al., 2019; Vilcinskas et al., 1997). There are also evidences that the fungal toxins destruxins (DTXs) alter circulating hemocytes of *G. mellonella* larvae (Vey et al., 2002; Vilcinskas et al., 1997). But in *Drosophila*, Destruxin does not appear to affect *Drosophila* cellular immune responses *in vivo* but would rather suppress the humoral immune response in *Drosophila* (Pal et al., 2007). Here, the use of *NimC1^1^; Eater^1^* did not reveal any major role of plasmatocytes in host defense against the five entomopathogenic fungi. Flies lacking these two receptors are unable to phagocyte bacteria, latex beads or apoptotic corpses (Bretscher et al., 2015; Dolgikh et al., 2025; Melcarne et al., 2019b) and have lost the ability to bind to *M. anisopliae* spores (Figs. S1E, F). The observation that hemoless flies lacking hemocytes due to the over-expression of *Bax* display a wild-type survival rate to *B. bassiana* also support the notion that the humoral response is more critical than cellular response to combat this entomopathogenic fungus (Figs. S1A, B). We did however observe an increased susceptibility of hemoless flies to *M. anisopliae* (Figs. S1C, D) pointing to a role of hemocytes against this species that could not be uncovered with *NimC1^1^; eater^1^* fly line. While the cellular response is important against yeast like *C. glabrata* (Quintin et al., 2013), filamentous fungi may evade phagocytosis by reducing the exposition of surface molecule(s) that are associated with non-self-recognition, as inferred in studies in other insects (Gillespie et al., 2000; Wang and St. Leger, 2006). Additional studies are required to better define the role of the cellular response to entomopathogenic fungi.

A key aim of our study was to compare *Drosophila* immune responses to fungi with distinct ecological strategies. Despite lifestyle differences, we found a largely conserved requirement for Toll signaling, antifungal peptides, and melanization across all species tested. However, notable differences emerged. The two generalist entomopathogenic species strongly activate the Toll pathway but can overcome Toll effectors to infect and kill flies. Studies have shown that these species express many virulence factors that likely confers them protection against the host immune system (Gao et al., 2011; Hu et al., 2014; Wang and Wang, 2017; Yuan et al., 2020). In contrast, *A. fumigatus* is weakly pathogenic upon natural infection and accordingly does not activate the Toll pathway. This opportunistic fungus does not have the ability to effectively cross the cuticle of *Drosophila*, reflecting an opportunistic lifestyle. In contrast the specialist noctuid-virulent fungus *M. rileyi* was moderately pathogenic to flies and only triggered the Toll pathway at a late time point. This may reflect the proliferation of this species within the host under a yeast-like blastospore form with a thinner cell wall (Boucias et al., 2016; Liu et al., 2019), that would be less recognized by the sensors acting upstream of the Toll pathway. This infectious strategy differs from the toxin-mediated pathogenesis seen in *B. bassiana* and *M. anisopliae* during fly colonization (Boucias et al., 2016; Schrank and Vainstein, 2010; Wang and Wang, 2017; Wang et al., 2021). Strikingly, we observed that *E. muscae* was greatly pathogenic to flies but did not trigger a strong Toll pathway response. Nevertheless, the Toll and melanization modules do contribute to host defense to *E. muscae* indicating that this species is also sensitive to immune effectors. Our study finds that *E. muscae* has developed a specific colonization strategy to escape the host immune response. After crossing the cuticle barrier and reaching the hemolymph, *E. muscae* grows as multinucleated protoplasts undetected by the immune system. The absence of cell wall in protoplasts results in little glucan release into the hemolymph, which might prevent recognition. This strategy would explain the high pathogenicity of *E. muscae,* allowing growth hidden from the immune system, and only differentiating into thinner cell-walled hyphae only shortly before the death of the host. While we cannot however rule out that the high pathogenicity of *E. muscae* may also be partly due to the fungus’s increased resilience to immune effectors, we favor the interpretation that it is instead mainly driven by its capacity to evade immune detection. Similar escape strategies to hide from the host immune response have been described in specialists such as *M. rileyi* and *O. sinensis* (Boucias et al., 2016; Li et al., 2020b; Liu et al., 2019). Thus, two distinct infection strategies emerge among entomopathogenic fungi: (1) Generalists, such as *B. bassiana* and *M. anisopliae*, mount an aggressive infection, are immediately recognized by the immune system, and rely on a broad array of virulence factors to counter host defenses (Gao et al., 2011; Schrank and Vainstein, 2010; Wang et al., 2021); and (2) Specialists, such as *E. muscae*, *M. rileyi*, and *O. sinensis*, adopt stealthier strategies, growing initially in immune-evasive forms before lately transitioning to lethal hyphal stages. These strategies appear to correlate with the fast and slow timing of host infection: for example, *O. sinensis* persists for months in *Thitarodes* larvae, only initiating pathogenic development later (Liu et al., 2019; Meng et al., 2015; Rao et al., 2019). Similarly, the slow-growing, behavior-manipulating *E. muscae* may rely on immune evasion to enable host manipulation and spore dispersal.

Taken together, our study clarifies the immune strategies employed by *D. melanogaster* to combat fungal infections. We further provide insight on how entomopathogenic fungi circumvent the immune response. Our study complements the recent identification of fungal virulence factors required to disarm the immune system (Lu et al., 2024; Shang et al., 2023; Tang et al., 2025). The interplay between local and systemic immune responses highlights the complexity of antifungal defense mechanisms in *Drosophila*. Understanding the evolutionary pressures that shape immune responses to fungal pathogens could have broader implications for controlling fungal infections in both natural and agricultural settings.

## Materials and methods

### Fly stocks

The fly stocks iso *w^1118^* drosdel wild-type was used as the genetic background for mutant isogenization (Ferreira et al., 2014). For survival experiments, *w^1118^ drosdel*, Oregon R or Canton S flies were used as wild-type controls. To determine the contribution of the four main immune modules to survive different fungal species, the *Relish* (*Rel^E20^*), *spätzle* (*spz^rm7^*), *PPO1* and *PPO2* double mutant (*PPO1^△^*, *PPO2^△^*), *NimC^1^* and *eater* double mutant (*Nim C1^1^; Eater^1^*) (Binggeli et al., 2014; Dudzic et al., 2019; Hedengren et al., 1999; Lemaitre et al., 1996; Melcarne et al., 2019a) were used. The fly line devoid of all four immune modules: Imd, Toll, Phagocytosis and Melanization [genotype: *Def(Hayan-psh)*; *NimC1; Eater*, *Relish^E20^*] (Ryckebusch et al., 2025) is used to investigate whether immune modules function independently or synergistically to combat different fungi.

To compare the importance of Toll sensing pathway, we used flies lacking Hayan (*Hayan^SK6^*), Psh (*psh^SK1^*), both Hayan and Psh (*Hayan-psh^def^*) (Dudzic et al., 2019), Gram-negative bacteria binding protein 3 (*GNBP3^hades^*) (Gottar et al., 2006), ModSP (*modSP^1^*), both Psh and ModSP (*psh^1^;;modSP^1^*) (Buchon et al., 2009).

To determine the role of host defense peptides to combat fungal infection, we used flies deleting ten of the twelve Bomanins (*Bom^△55C^*) (Clemmons et al., 2015) and flies lacking Metchnikowin (*Mtk^R1^*), Drosomycin (*Drs^R1^*) (Hanson et al., 2019), Baramicin (*BaraA^△^*) (Hanson et al., 2021),Daisho 1, 2 (*△Dso*) (Cohen et al., 2020), as well as compound mutants *Mtk^R1^*;*Drs^R1^*, *△AMP10* and *△AMP14* (Carboni et al., 2022; Hanson et al., 2019). In this study, we generated four new compounds mutants lack host defense peptides in the Drosdel background, (*BaraA^SW1^*, *△Dso*); (*Mtk^R^*, *BaraA^SW11^*; *Drs^R1^*); (*Mtk^R^*, *BaraA^SW1^, △Dso ^1^*; *Drs^R1^*) and (*Mtk ^R1^*, *Bom^△55C^*; *Drs^R1^*). The *Hml Δ-GAL4, UAS-GFP* fly line and UAS-Bax were used to delete hemocytes as described in (Defaye et al., 2009; Gaumer et al., 2000). We used a newly generated *w;; Hml^P2A^-GAL4, UAS-CaMPARI2* to monitor hemocyte number and localization upon fungal infection (Moeyaert et al., 2018; Stephenson et al., 2022) The wild-type and mutant flies used in this study are listed in Table S1.

### Microbial cultures

The wild type strains of *B. bassiana* 802, *M. anisopliae* ARSEF 2575, *M. rileyi* PHP1705 and the genetic strains *M. anisopliae* tagged GFP were cultured on malt agar (1g Bacteriological Peptone, 20g Glucose, 20 g Malt Extract, 15 g Agar, with the addition of water to the total volume of 1 liter) at 29℃ for two weeks for harvesting conidial spores. The wild type strain *A. fumigatus* was cultured on malt agar at 37℃ for one week for harvesting spores. Fungal materials were then strained through three layers of precision wipes with sterile water to collect spores, which were resuspended with 0.05% Tween 80 before being used for infection. The *E. muscae* ARSEF 454 was cultured in Grace’s Insect Medium supplemented with 10% fetal bovine serum at 21℃. A liquid of 1 ml growing cultures after three weeks culture was transferred to a new cell culture flask with the addition of 10 ml media to maintain the fungal strains *in vitro*. The Gram-positive bacteria *Micrococcus luteus* were cultured on Luria Broth at 37℃ and the optical density of the pellet (O.D.) at 600 nm 200 were used for infection.

### Survival experiments

Survival experiments were conducted by both topical infection (natural infection) and systemic infection (septic infection), with twenty 2- to 4-day-old adult males per vial with two to three replicate experiments. Topical infection assays were performed by immersing flies in spore suspensions for 30 s. The following concentrations (in spores/ml) of spore suspensions were used: *A. fumigatus, B. bassiana* and *M. rileyi*: 1 × 10^9^; *M. anisopliae*: 5 × 10^8^. For *E. muscae* natural infection, about twenty-five 1- to 2-day-old adult male flies were infected by spore shower from twenty freshly emerged cadavers that were previously injected with protoplasts for 24 h. The average spores ejected from each female or male fly for 24 h were evaluated as described in (Edwards and De Fine Licht, 2024). Briefly, the head of the cadaver was insert into Vaseline inside the lid of the 2 ml tube containing 1ml 1% Triton-X and 0.2% maleic acid solution for a period 24 h to collect ejected of spores. Then, spores in solution were counted on a hemocytometer under a microscope. Systemic infections were performed by pricking 2- to 4- day-old adult males in the thorax with a 100 µm-thick insect pin dipped into spore suspensions. The following concentrations (in spores/ml) of spore suspensions were used: *B. bassiana, M. anisopliae* and *M. rileyi*: 1× 10^8^, *A. fumigatus*: 1 × 10^9^. Infected flies were flipped at the end of the first day and maintained at 29℃ and flipped every two days later. Systemic infection with *E. muscae* was performed by injecting 23 nl of protoplast suspension per fly with a concentration of 2× 10^6^ protoplasts/ml in PBS. The mortality was recorded daily. At least three independent experiments for survival to infection were performed with 20 flies per vial on standard fly medium.

### Gene expression levels by RT-qPCR

To investigate the immune responses to fungal infections, we performed the quantitative reverse transcription PCR (qRT-PCR) analysis of Drs and BomBc3 used as read-outs of Toll pathway activity. About 10 adult flies were collected at indicated time points and homogenized for RNA extraction using TRIzol reagent. The total RNA was resuspended in RNase-free water and 0.5 mg of total RNA in 10 ml reactions was reverse transcribed using PrimeScript RT (TAKARA) with random hexamer and oligo dT primers. Quantitative PCR was performed on a LightCycler 480 (Roche) in 96-well plates using PowerUp SYBR Green Master Mix. Primers were listed in Table S2. The expression of *Drs* was also investigated by observing GFP intensity of *Drs-GFP* flies after fungal infection (Ferrandon, 1998). Infected flies were anesthetized with CO_2_ and observed for fluorescence with a Leica Application SuiteX and LEICA DFC 7000T camera.

### Quantitation of fungal loads for kinetic growth

Fungal load in infected flies was measured either by qPCR or by fluorescence intensity of *M. anisopliae t*agged GFP. Adult male flies infect with *B. bassiana* and *M. anisopliae* were incubated at 29°C and *E. muscae* at 21°C. Groups of 10 flies were sampled at indicated time points and frozen in liquid nitrogen. The total DNA of each sample was extracted with DNeasy Plant Mini Kit (Qiagen). Fungal loads of *B. bassiana* and *E. muscae* was quantified by fungal ribosomal protein 18S rRNA relative to the *Drosophila* ribosomal protein Rpl32 (Rp49), while fungal load of *M. anisopliae* was quantified by fungal ribosomal protein Rpl32 relative to *Drosophila* protein Actin. Primers were listed in Table S2. *M. anisopliae* tagged GFP was also used to investigate the growth of fungi inside the flies after pricking. The infected flies were anesthesized with CO_2_ and observed for fluorescence with a Leica Application SuiteX and LEICA DFC 7000T camera.

### Ex-vivo phagocytosis assay with microscopy

Five third-instar larvae were cleaned and dissected in 150 µL of Schneider’s medium, ensuring that vital organs remained intact. The samples were incubated with approximately 10⁵ bacterial bioparticles or fungal blastospores in a humidified chamber for 2 hours. Cells were then fixed with 4% paraformaldehyde (PFA) and stained with phalloidin (1:200) for 1 hour at room temperature, followed by three washes with 1× PBS. Finally, slides were mounted with a mounting medium and sealed with nail polish. Hemocytes were imaged using an inverted Evident confocal microscope equipped with a 60× oil immersion lens. Maximum Z-projections were generated for analysis using FIJI/ImageJ software.

### Ex-vivo phagocytosis assay with FACS

Five third-instar larvae were surface-sterilized and bled into 130 µL of Schneider’s medium within 2 minutes. The hemolymph was collected in Protein Lo-Bind Eppendorf tubes to minimize sample loss. Approximately 10⁵ bacterial bioparticles or fungal blastospores were added to the hemocyte suspension and incubated at room temperature for 2 hours. Following incubation, the samples were placed on ice, and hemocytes were immediately analyzed using a CytoFLEX flow cytometer (Beckman Coulter) to measure the fraction of cells phagocytosing. 60 μl volume was read at medium speed (40 μl/min).

Hemocytes from Hml GAL4, UAS-GFP larvae were used to define the hemocyte population for flow cytometry analysis. GFP-positive cells were identified and gated based on fluorescence detected in the B525 channel, enabling the exclusion of debris and non-hemocytic events. The same gating strategy was applied consistently across all genotypes to track the hemocyte population and to quantify the proportion of hemocytes exhibiting fluorescence corresponding to internalized bacterial bioparticles or fungal spores. The percentage of hemocytes phagocytosing (*f*) was calculated using the following equations:

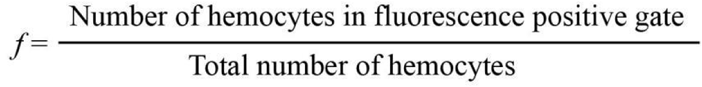

## Statistical analysis

Statistical analyses were performed using GraphPad Prism 10 software. Survival experiments were repeated at least three times independently with 20 flies per treatment. Statistical significance of survival data was calculated with a log-rank test (Mantel-Cox test) comparing each genotype to *w^1118^*flies. Quantitative PCR data and fungal load data were compared by One-way ANOVA following Tukey’s multiple comparisons test. The full statistic models and outputs of Figures are listed in Table S3. P values of < 0.05 = *, < 0.01 = **, < 0.001 = ***, and < 0.0001 = **** were considered significant. ns indicates non-significant, P>0.05. Error bars represent the standard deviation (s.d.) of replicate experiments.

## Acknowledgments

We thank Samuel Rommelaere, Jean-Philippe Boquette, Fanny Schüpfer for experimental help. We thank Kathryn Bushley and Carolyn Elya for providing the *Entomophthora muscae* strains used for this study. Guiqing Liu stay in the Lemaitre lab was supported by funding from the China Scholarship Council (CSC, No. 202208440111) and the SNSF grant TMAG-3_225935. We thank Vienna Drosophila Resource Center (VDRC) and the Bloomington stock center for fly stocks. This project was supported by the SNSF grants (310030_215073).

## Author Contributions

Conceived and designed the experiments: GQL BL. Performed the experiments: GQL YT JL PKS. Analyzed the data: GQL YT PKS BL. Contributed reagents/ materials/analysis tools: MAH BL. Wrote the paper: GQL MAH BL.

## Supplementary Figure Legends

**Figure S1:**
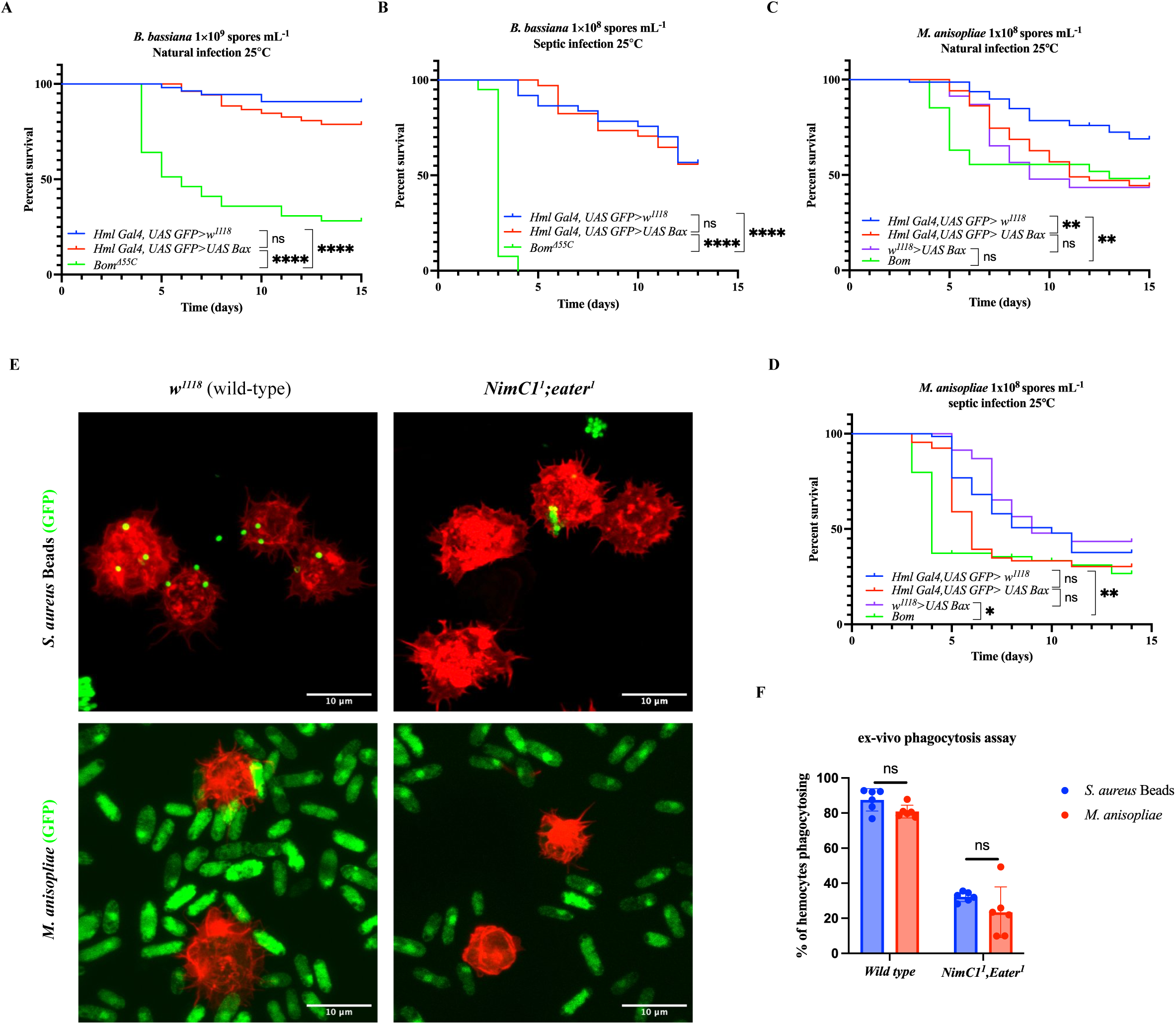
Contribution of the cellular response to host defense to fungal infection. (A-D) Survival analysis of males lacking most plasmatocytes—due to Bax overexpression in plasmatocytes (*HmlΔ-GAL4, UAS-GFP / UAS-Bax*) (Defaye et al., 2009)—shows only a modest reduction in their ability to survive natural or septic injury infection with *B. bassiana* and *M. anisopliae*. Given the numerous roles of hemocytes in immunity and metabolism (Westlake et al., 2024b), these experiments cannot disentangle the specific contribution of hemocytes to antifungal defense. Both systemic and natural infections were carried out at 25 °C. *Bom^Δ55C^* flies were included as a positive control. (**E**). Spores of *M. anisopliae-GFP* bind to wild-type but not *NimC1^1^;eater^1^* deficient plasmatocyte (stained in red with phalloidin). Plasmatocytes from third instar larvae were incubated in presence of *M. anisopliae-GFP* for 2 hours. Representative image are shown. (**F**) *NimC1^1^;eater^1^* deficient plasmatocytes display reduced ability to phagocyte or bind to spores of *M. anisopliae.* Plasmatocytes from third instar larvae were incubated in presence of *M. anisopliae-GFP* or bead of *S. aureus* for 2 hours. Note that this assay does not allow to distinguish if plasmatocytes bind to spores or uptake it internally.

**Figure S2.**
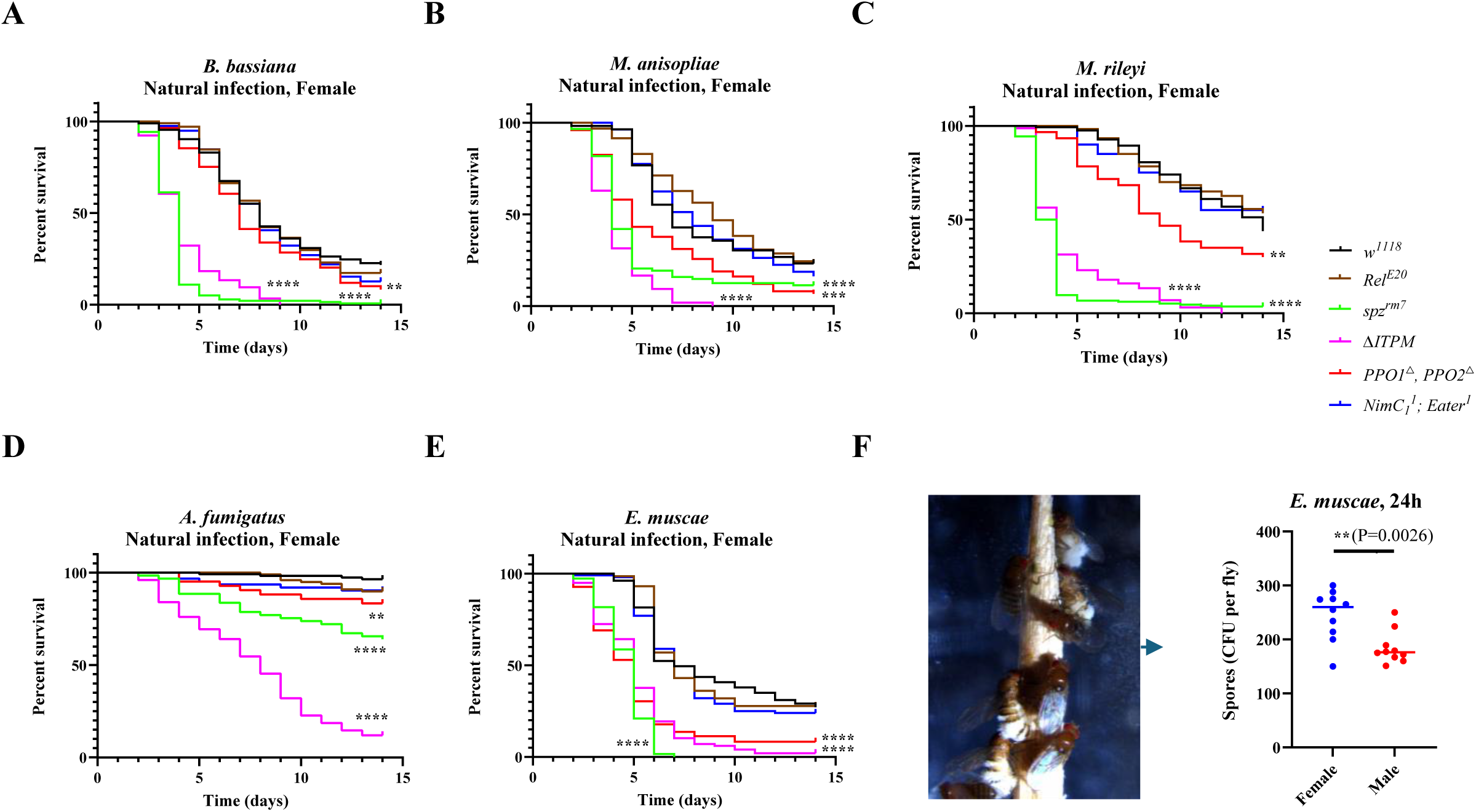
Survival of the female wild-type and mutant flies deficient for the four main immune modules upon natural infection with the five fungi. (**A-E**) Survival rate of the females shows that *spz^rm7^* flies (P<0.0001) and *PPO1, PPO2* double mutant flies (P<0.0001) are severely affected in their capacity to survive infection with the five fungi compared to wild-type *w^1118^* flies. (**F**) The evaluation of spore concentration after the 24 h ejection from the fresh cadavers shows that more spores were produced from female cadavers than male cadavers (**, P<0.01). For natural infection of *E. muscae*, a total of 10 female and 10 male fresh cadavers were used for one infection treatment by spore shower. Data were analyzed using the Log rank test and values are pooled data from at least three independent experiments. Full statistical details are available on Table S3. Related to Figure 1.

**Figure S3.**
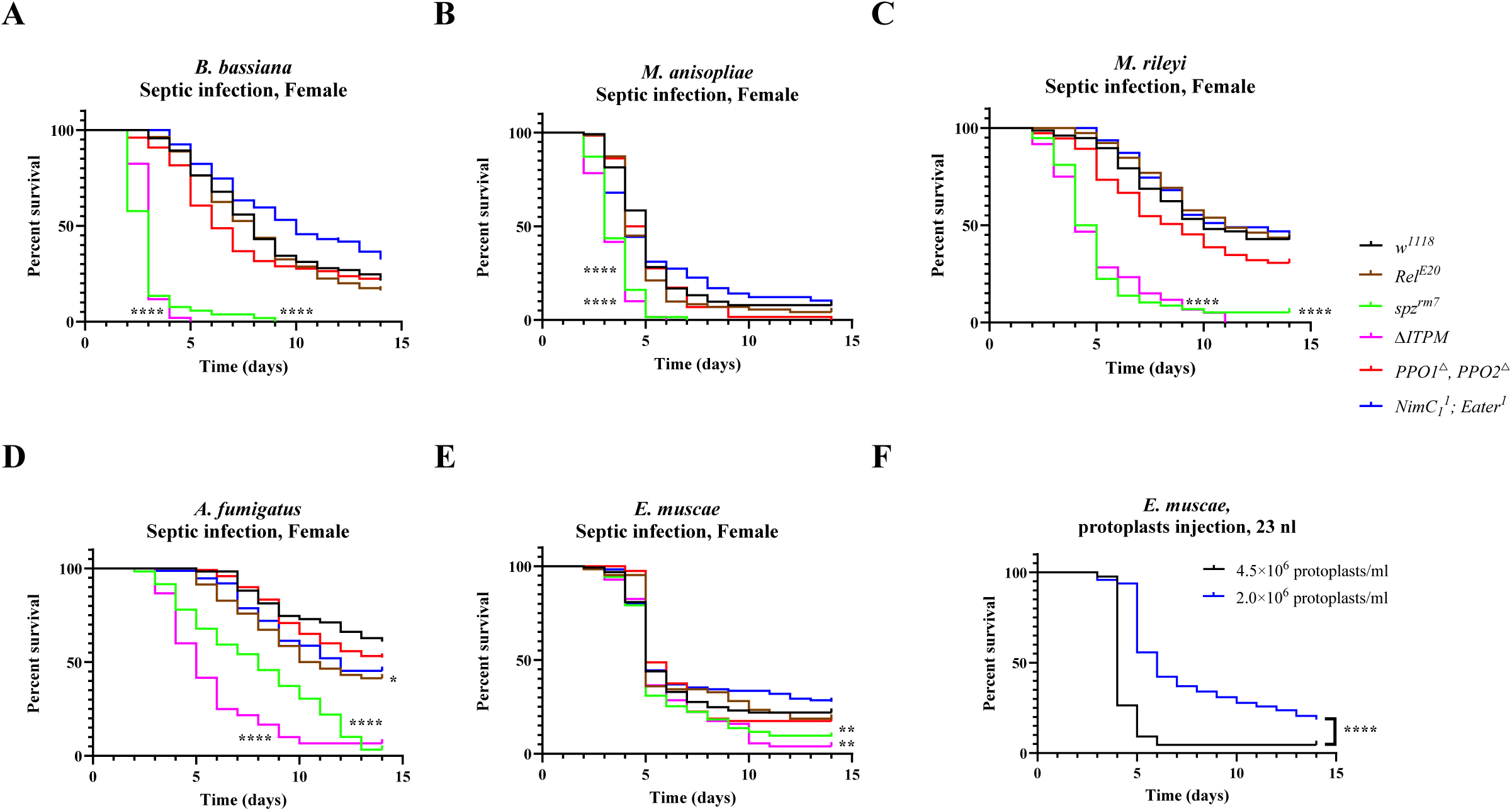
Survival of the female wild-type and mutants devoid of the four main immune modules upon septic infection with the five fungi. (**A-C**) Survival rate of females following pricking with spore suspensions show that *spz^rm7^* flies (P<0.0001) contribute most to surviving infection by the *B. bassiana*, *M. anisopliae*, and *M. rileyi*. (**D**) *spz^rm7^* and to lower extent *Rel^E20^* females exhibited a strong susceptibility to infection with *A. fumigatus* (P<0.0001). (**E**) No significant difference in survival of the four main modules upon infection with *E. muscae* by the injection of protoplasts compared to wild-type females. (**F**) Survival of *w^1118^* flies upon infection with *E. muscae* showed that faster killing was found when injected with a higher concentration of protoplasts (****, P<0.0001). Data were analyzed using the Log rank test and values are pooled data from at least three independent experiments. Full statistical details are available on Table S3. Related to Figure 2.

**Figure S4.**
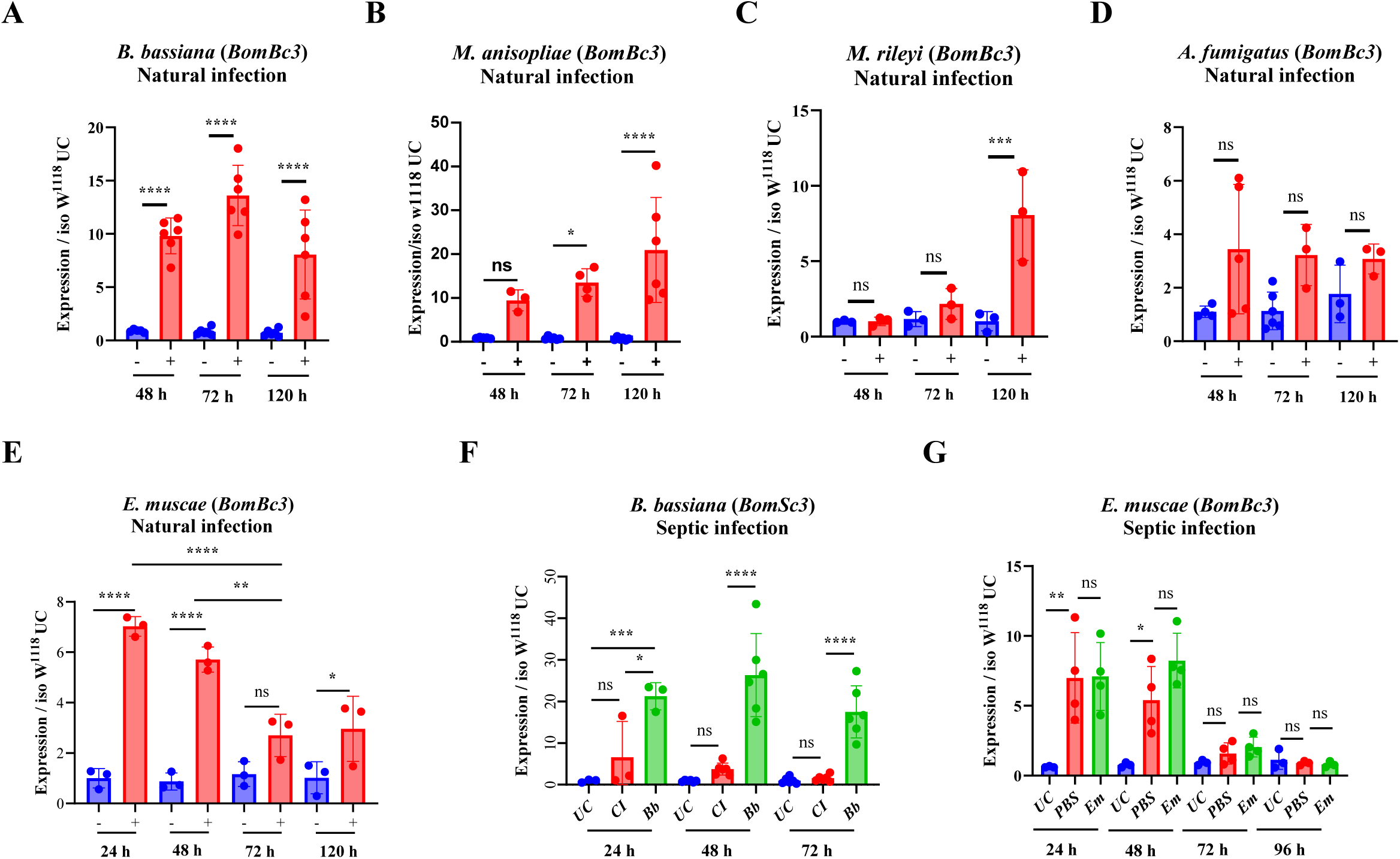
Expression of Bomanin (*BomBc3*) in wild-type flies upon infection with the five fungi. (**A-C**) Natural infection with *M. anisopliae* induced a highest expression of *BomBc3,* followed by *B. bassiana* and *M. rileyi* only induced the expression of *BomBc3* at a late stage, at 120 h after infection. (**D**) Natural infection with *A.fumigatus* could not induce the expression of *BomBc3.* (**E**) Natural infection with *E. muscae* induced a low expression of *BomBc3* at 24 h and 48 h after infection, then followed by a significant decrease. (**F**) Septic infection with *B. bassiana* spores activates higher *BomBc3* expression compared to natural infection and clean injury with a needle. (**G**) Expression of *BomBc3* after injection with *E. muscae* protoplasts was similar to the injection with PBS, indicating protoplasts cannot activate Toll signaling. Expression was normalized with *w^1118^* UC set as a value of 1. Data were analyzed using One-Way ANOVA followed by Tukey’s multiple comparison tests. ns, P>0.05; *, P<0.05; **, P<0.01; ***, P<0.001; ****, P<0.0001. Values represent the mean ± s.d. of at least three independent experiments. Full statistical details are available on Table S3. Related to Figure 3.

**Figure S5.**
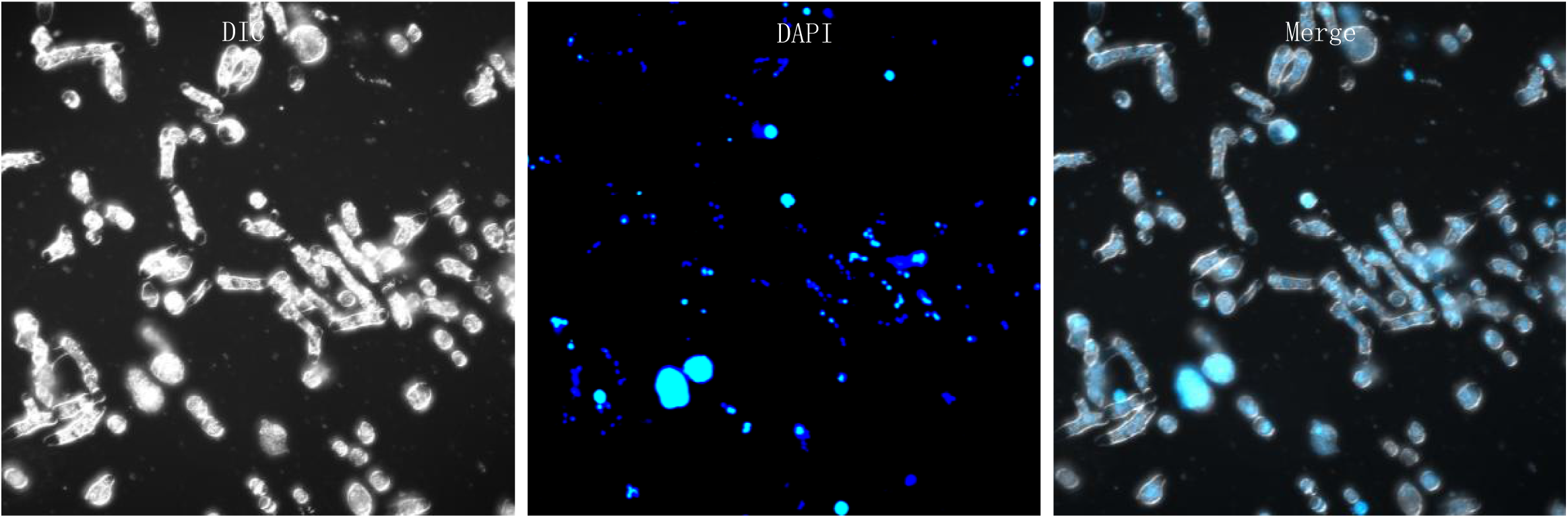
DAPI staining of the *E. muscae* protoplasts.

**Figure S6.**
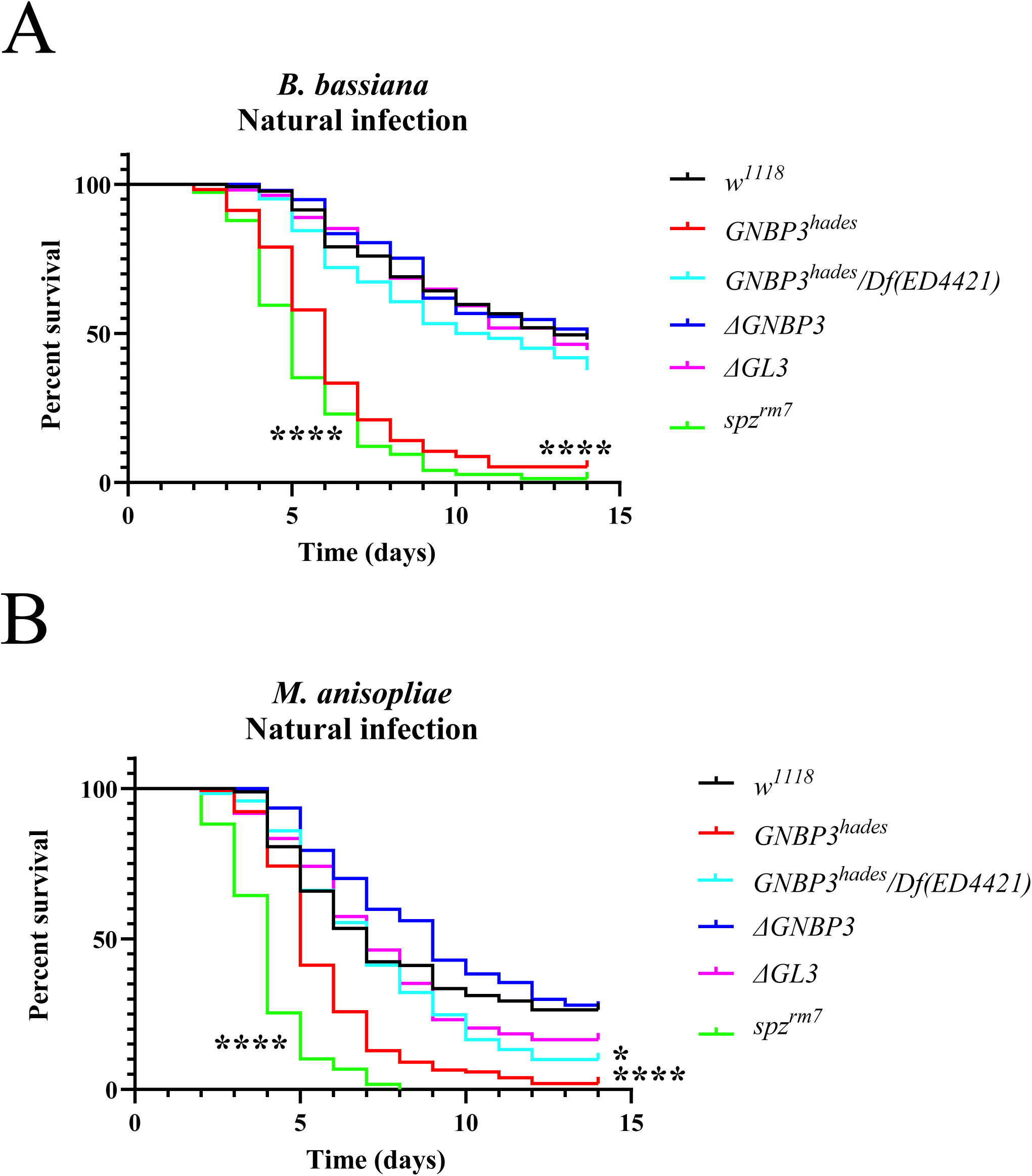
Contribution of GNBP3 and GNGP like 3 to survival upon natural infection with *B. bassiana* and *M. anisopliae*. (**A-B**) Survival rate of male flies following infection with the two fungi shows that both GNBP3 and GNGP like 3 mutant flies are not susceptible or slightly susceptible to *B. bassiana* and *M. anisopliae*. Full statistical details are available on Table S3. Related to Figure 5.

**Figure S7.**
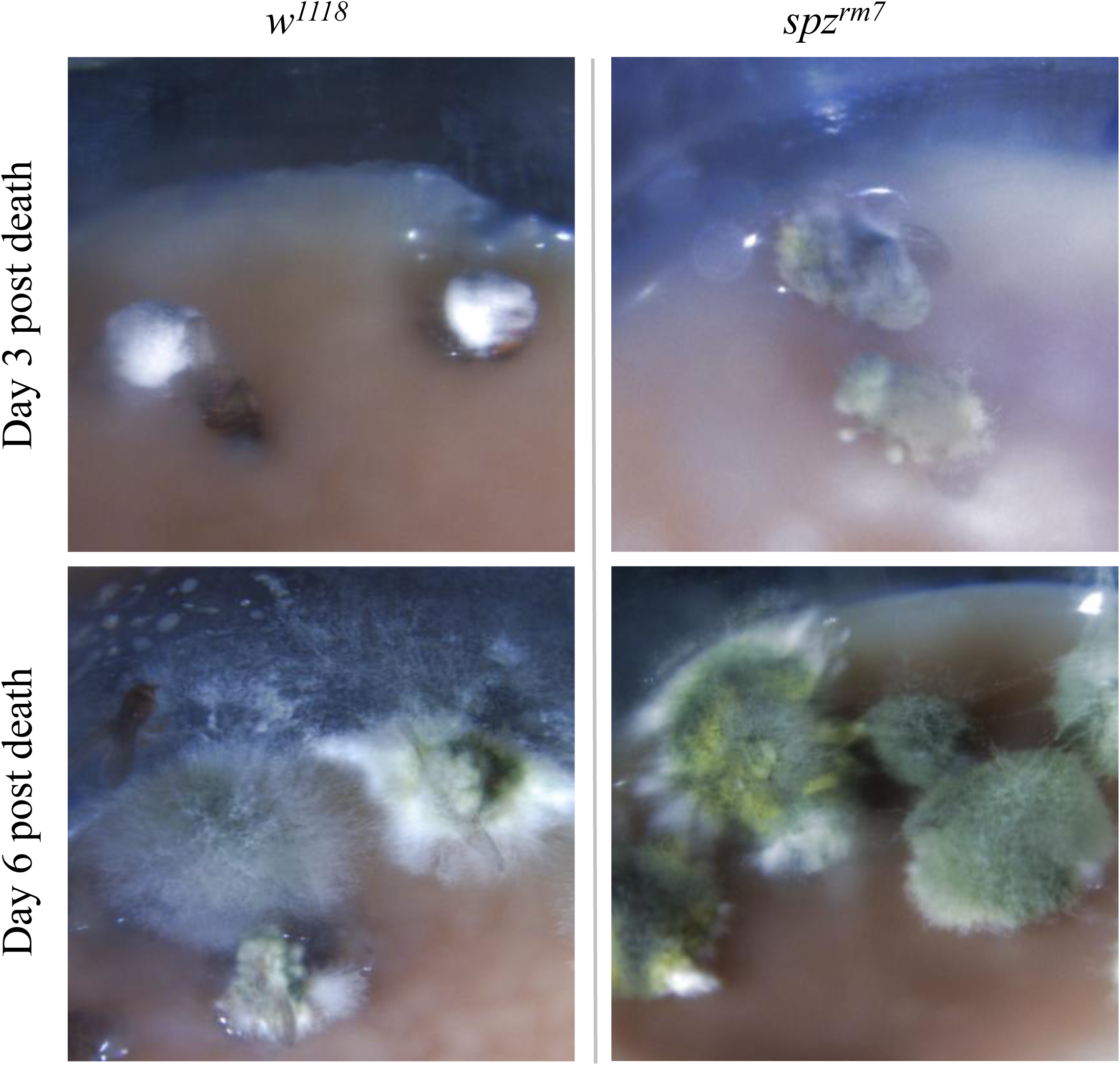
Fungal dissemination from wild-type and *spz^rm7^* cadavers after natural infection with *M. anisopliae*. *Metarhizium anisopliae* extruding from cuticle and growing in *spz^rm7^* cadavers are faster than that in wild-type cadavers. *M. anisopliae* can eventually disseminate through the whole flies both in wild-type and *spz^rm7^* cadavers. Related to Figure 6.

**Figure S8.**
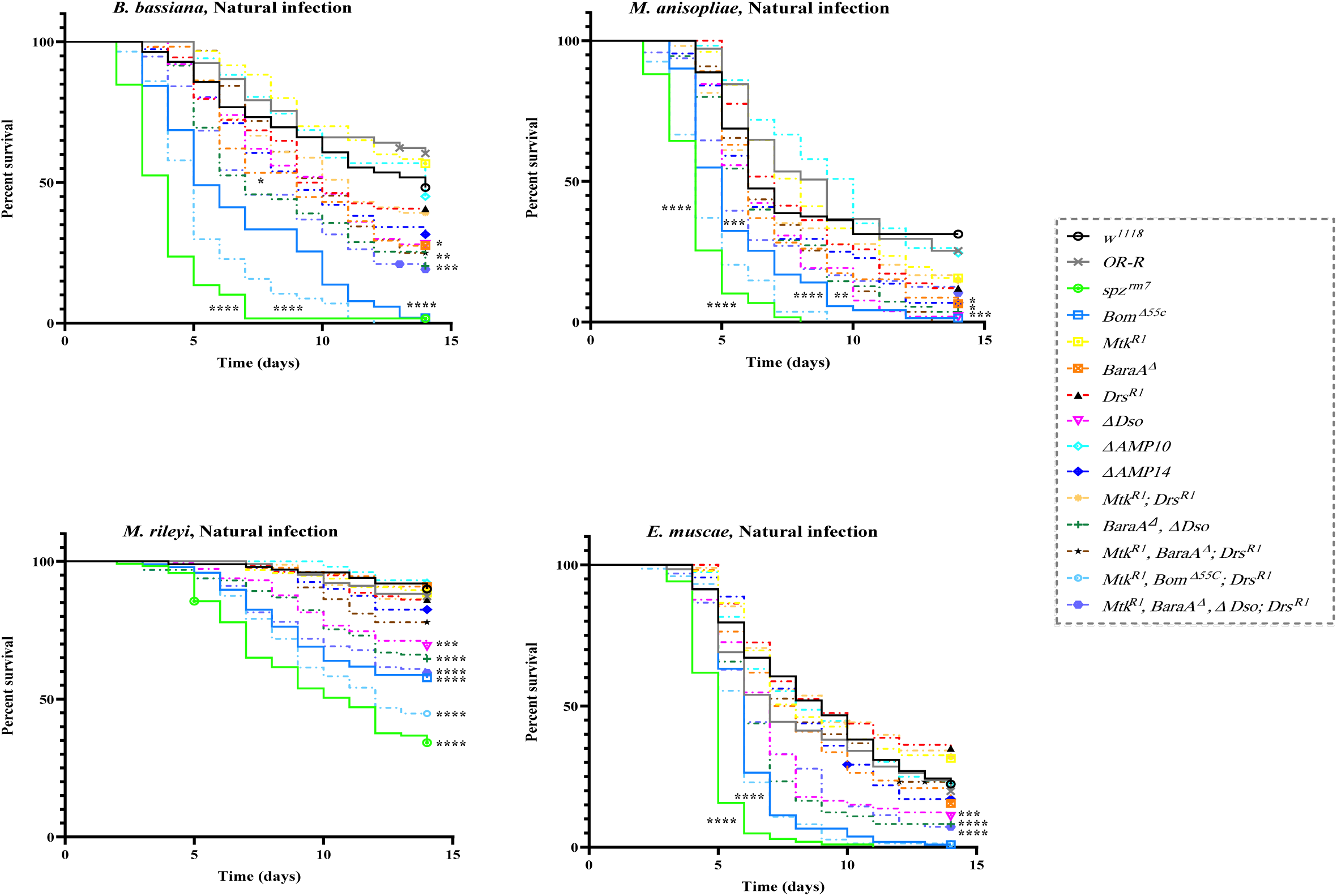
Sum survival curves of single and compound host defense peptides mutants after infection with *B. bassiana*, *M. anisopliae*, *M. rileyi* and *E. muscae*. Full statistical details are available on Table S3. Related to Figure 7.

**Figure S9.**
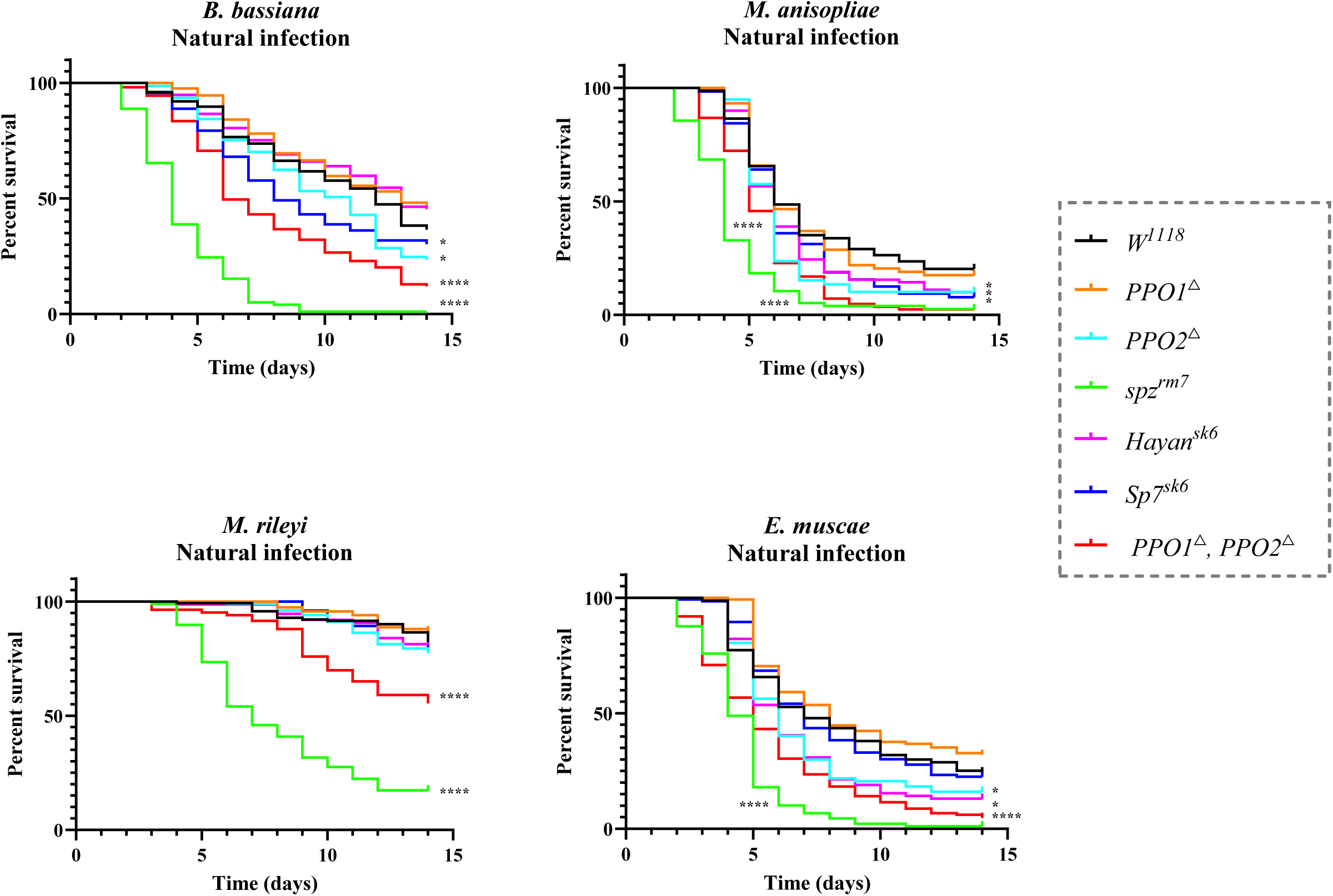
Sum survival curves of mutants of SPs involved in melanization after infection with *B. bassiana*, *M. anisopliae*, *M. rileyi* and *E. muscae*. Full statistical details are available on Table S3. Related to Figure 7.

**Figure S10.**
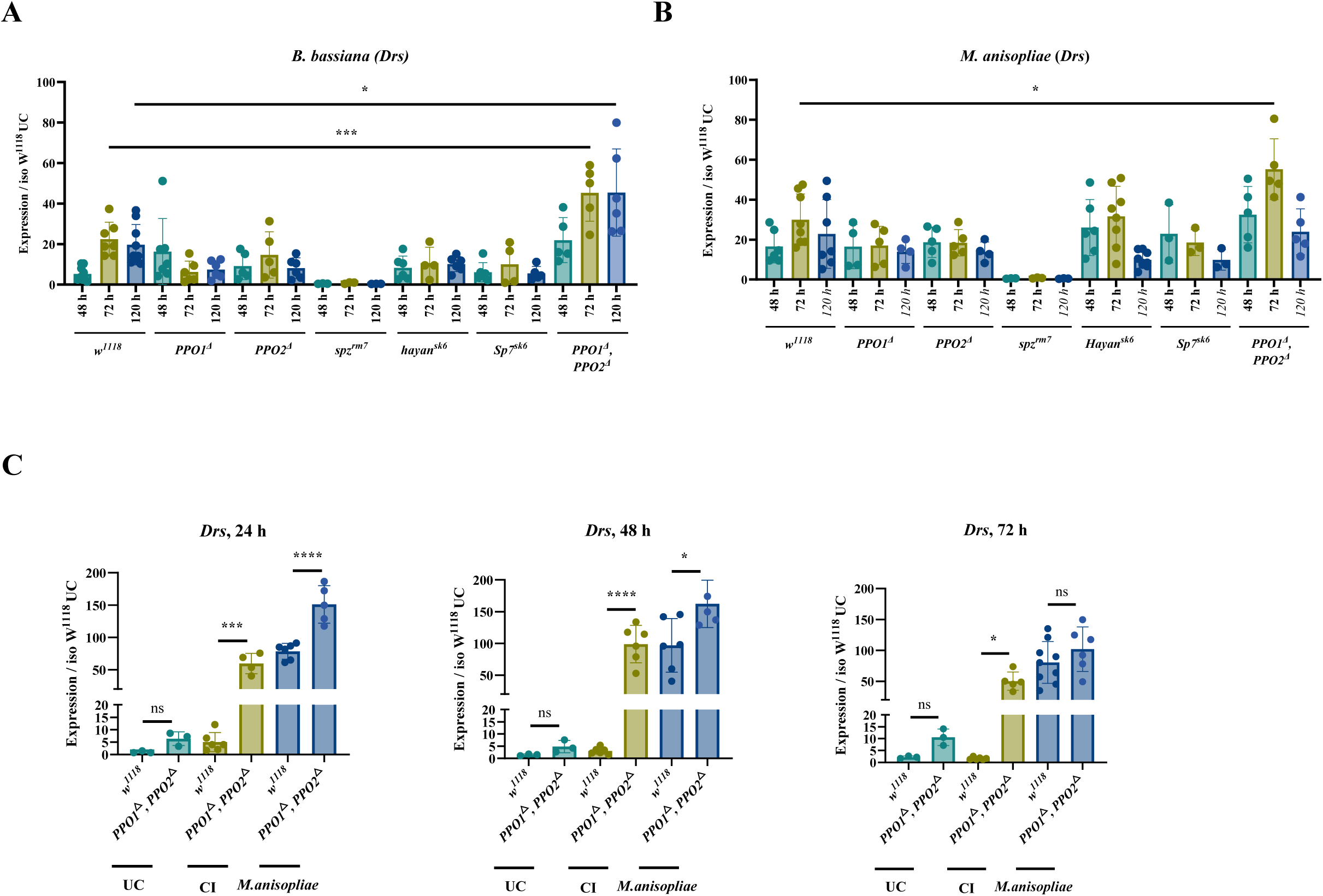
Expression of *Drosomycin* of mutants of SPs involved in melanization upon natural and septic infection with *B. bassiana* and *M. anisopliae*. (**A-B**) Statistical analysis reveals higher *Drosomycin* expression in *PPO1, PPO2* double mutant compared to wild-type at 72 h and 120 h after natural infection with *B. bassiana* and at 72 h after natural infection with *M. anisopliae* (*, P<0.05; ***, P<0.001). The *spz^rm7^* flies as a positive control. (**C**) Compared with wild-type flies, PPO1, PPO2 double mutant flies exhibit an enhanced Toll pathway activity upon clean injury and septic infection with *M. anisopliae* (ns, P>0.05; *, p<0.05; ***, P<0.001; ****, P<0.0001). Expression was normalized with *w^1118^* UC set as a value of 1. Data were analyzed using One-Way ANOVA followed by Tukey’s multiple comparison tests and values represent the mean ± s.d. of at least three independent experiments. Full statistical details are available on Table S3. Related to Figure 7.

---

**Table S1.**
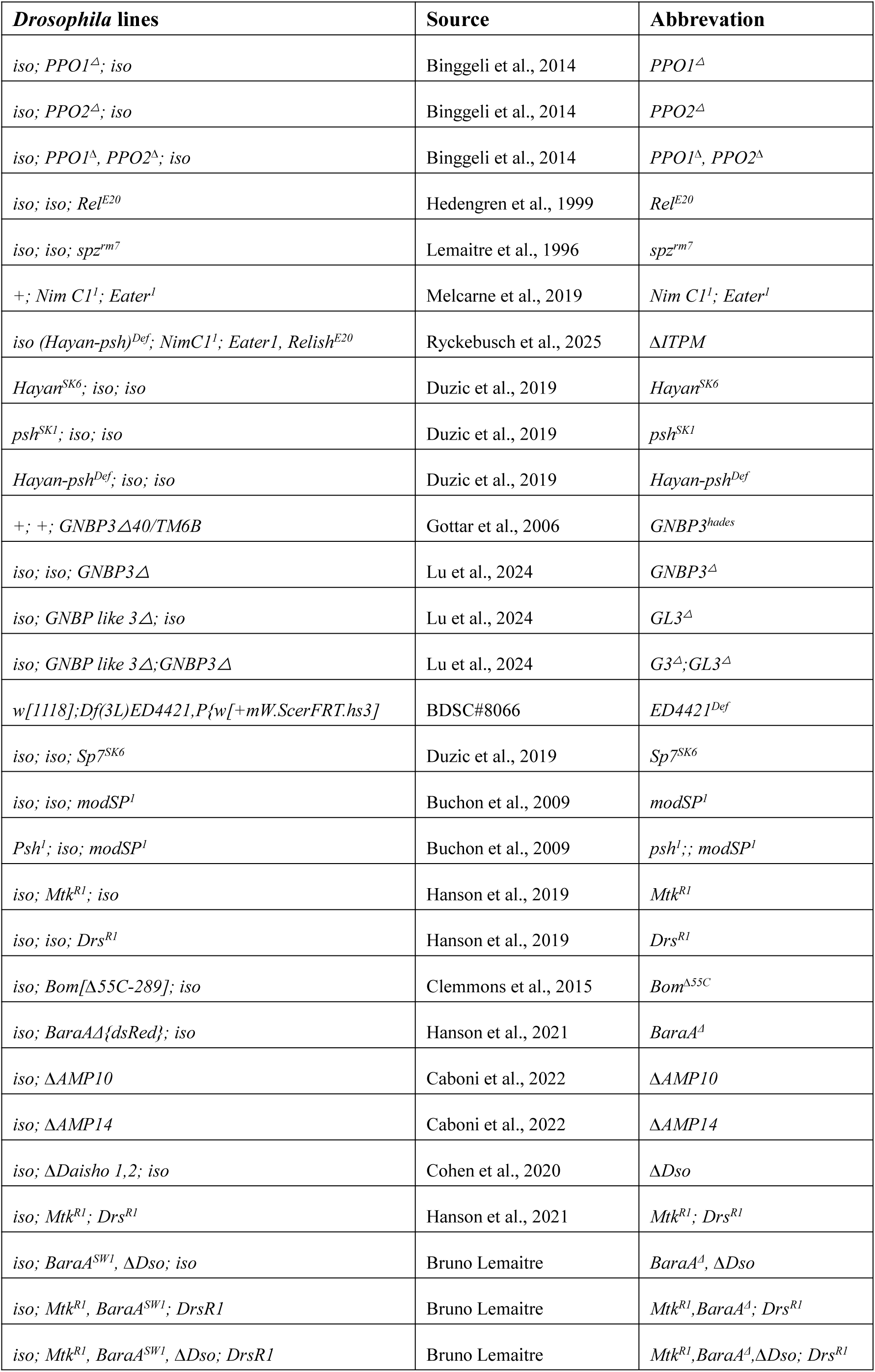
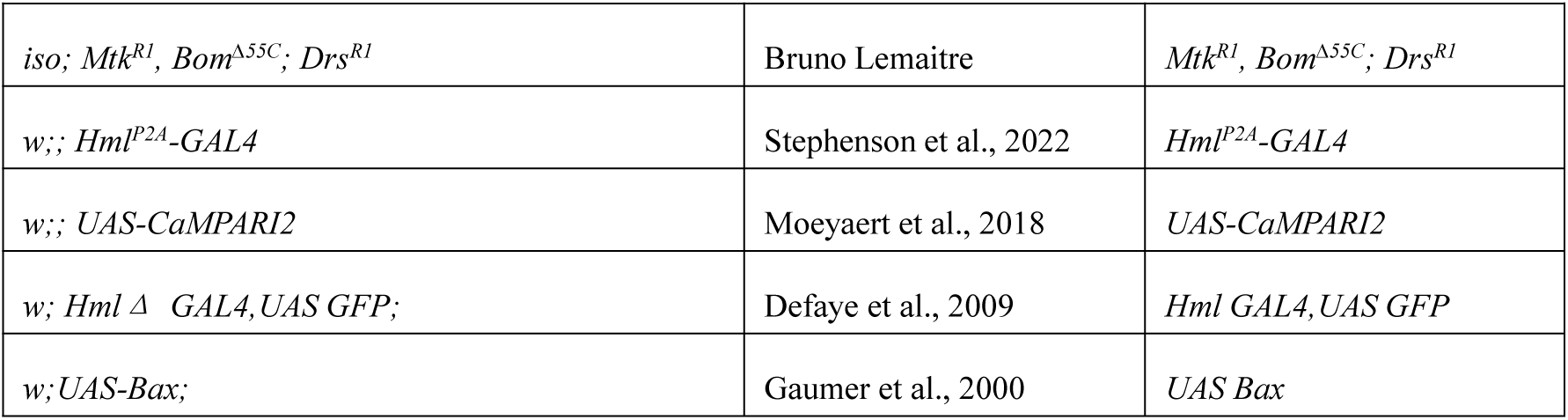
List of wild-type and mutant flies used in this study.

**Table S2.**
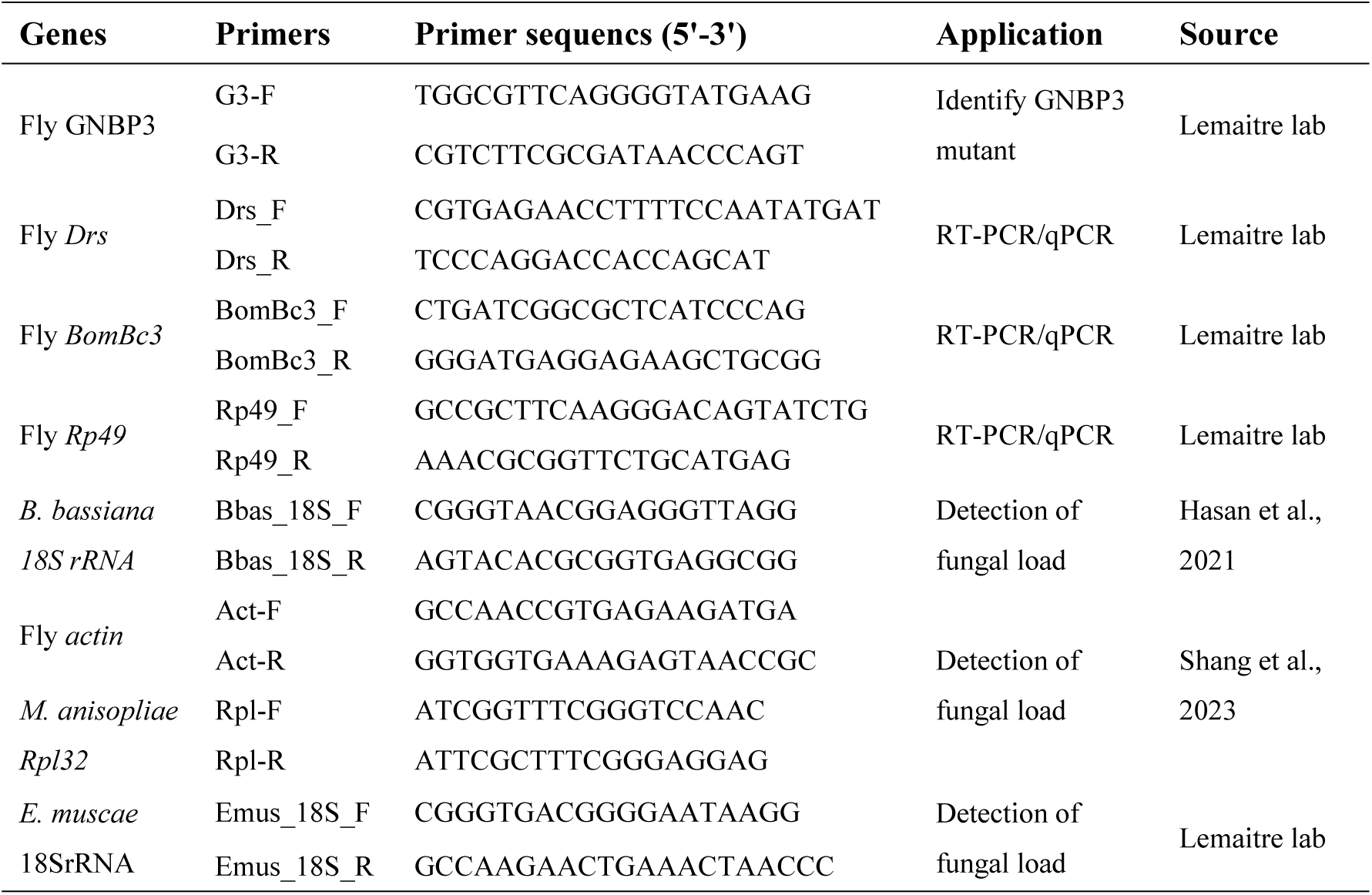
Primers used in this study.

